# Bridging scale-up to transplantation: pluripotent stem cell-derived pancreatic islet encapsulation in emulsion-generated high concentration alginate beads

**DOI:** 10.64898/2026.09.12.751148

**Authors:** Arianna Castro Rojas, Jonathan A. Brassard, Marie Billaud, Florent Lemaire, Robert Chen, Hamid Ebrahimi Orimi, Jiyu Jessica Tian, Steven Paraskevas, Corinne A. Hoesli

## Abstract

Pluripotent stem cell-derived pancreatic islet-like cell clusters (SC-islets) have emerged as potential cellular therapy for type 1 diabetes. While SC-islets can theoretically be produced in bioreactors to meet transplantation needs of many recipients per batch, biomanufacturing and translational challenges remain. SC-islet can be cultured in suspension using stirred tank or vertical wheel bioreactors, but these can impart hydrodynamic damage and lead to cellular agglomeration, particularly upon scale-up. We previously described a robust emulsion-based process to encapsulate murine pancreatic beta cells in high-concentration alginate beads which improved graft survival in allogeneic recipients. Here, we present a full pipeline for scalable microencapsulated SC-islet biomanufacturing with extended *in vitro* bioreactor culture leading into transplantation. We hypothesized that encapsulation would prevent cellular agglomeration, reduce mechanical stress, and preserve differentiation potential during scale-up. Encapsulation of Stage 6 SC-islets prevented cellular agglomeration during extended suspension culture (25 days) and increased cell recovery (91 ± 3%) compared with non-encapsulated aggregates (60 ± 10%). No significant differences in glucose-stimulated insulin secretion were observed with vs without microencapsulation. Stage 7 SC-islets matured in the bioreactor were transplanted either as encapsulated islets via the intraperitoneal route or as free SC-islets under the kidney capsule, where they displayed glucose-responsive human C-peptide secretion and remained functional *in vivo* for up to 98 days. Overall, this work establishes a scalable, robust, transplantation-ready encapsulation platform that supports SC-islet maturation and delivery, providing a generalizable strategy for scaling and transplanting encapsulated organoid systems.

## 1. Introduction

Type 1 diabetes is an autoimmune disease that leads to the destruction of insulin-producing pancreatic beta cells^1^. Islet transplantation has emerged as an alternative to daily insulin injections to regulate blood glucose levels. However, limited islet sources and the need for life-long immunosuppression to prevent graft rejection hinder widespread application. Stem cell–derived islets (SC-islets) represent a potentially scalable and renewable cell source for beta cell replacement therapy^2^. Early-phase clinical data indicate that hepatic SC-islet grafts can provide sustained reduction and even independence from exogenous insulin in individuals with type 1 diabetes placed under immunosuppression^3,4^.

Islet encapsulation creates a perm-selective barrier between the graft and components of the recipient immune system with the aim of reducing or eliminating the need from chronic immunosuppression^5,6^. Microencapsulated islets can survive for months to years in immunocompetent recipients, re-establishing glycemic control in rodent and primate diabetic animal models^7,8^. Past and ongoing clinical trials are evaluating the potential of this strategy in individuals with type 1 diabetes^2^. Moreover, encapsulation is expected to improve the safety of islet transplantation by allowing graft containment and potential retrieval if concerns arise. ^9^ Microencapsulation strategies have been explored across a range of bioprocessing applications to improve cell handling, including protection from hydrodynamic stress, control of cell aggregation, and facilitation of cell retention in perfusion systems. While these approaches have been extensively applied to microbial systems and certain mammalian cell cultures, their implementation in the context of pluripotent stem cell (PSC)-derived products remains limited^10–12^. The translation of microencapsulation strategies to complex, multicellular aggregates such as SC-islets raises additional challenges related to mass transfer, aggregate integrity, and functional maturation. The three-dimensional (3D) environment created by the encapsulation material also provides various mechanical stimuli that can impact differentiation outcomes^9^. Recent technological developments, including core–shell encapsulation platforms, have demonstrated the feasibility of high-throughput cell encapsulation^13,14^. However, these systems often rely on low-viscosity alginate formulations that typically are permeable to antibodies or lead to teardrop-shaped microbeads^13^ which may be problematic for transplantation applications.

We previously established a scalable, simple, robust and cost-efficient emulsion-based encapsulation method to encapsulate dispersed pancreatic beta cells^15^. We hypothesized that this process would permit encapsulation of cell clusters such as SC-islets, and that alginate immobilization would facilitate bioprocess scale-up by shielding SC-islets from hydrodynamic stress. After examining the impact of different alginate concentrations on SC-islet diameter, recovery and function after extended (25-day) suspension culture, we transplanted SC-islets in 5% alginate, previously shown to provide protection of beta cells in immunocompetent allogeneic mice^16^. We propose stirred emulsification and internal gelation as an accessible method to streamline SC-islet culture scale-up and transplantation in microbeads.

## 1. Materials and Methods

### 1.1. Cell source and maintenance

Human ESCs (WA01 H1, WiCell) were used under Canadian Stem Cell Oversight Committee and institutional review board (20-04-021, "Studying the differentiation of Pluripotent Stem Cells into pancreatic, vascular and other cells for cell therapy applications using 3D cultures and other bioreactors/bioprocessing") approval. The H1 cell bank was created at passage 25-30. Cells were frozen using mTeSR™1 medium (STEMCELL Technologies, 85850) with 10% Dimethyl sulfoxide (DMSO) (Fisher Scientific, BP231-100). Following thawing, the cells were cultured on Sarstedt (red) plates previously coated for 120 min with hESC-qualified Matrigel (Corning™, 354277) using 0.21 mL/cm^2^ mTeSR™1 medium (STEMCELL Technologies, 85850). Upon reaching 70-80% confluency, cultures underwent clump passaging via: (1) washing with Ca^2+^ and Mg^2+^ free DPBS (Gibco™, 14190144), (2) 4 incubation in 0.5 mM UltraPure™ Ethylenediaminetetraacetic acid (EDTA) diluted in Ca^2+^ and Mg^2+^ free Dulbecco’s phosphate-buffered saline (DPBS, pH 8.0, Invitrogen™, 15575020), (3) removing the EDTA solution and adding mTeSR™1 media (4) creating clumps using a cell scraper (Sarstedt, 83.3950), (5) gently triturating with a 10 mL pipette to obtain smaller clumps, (6) reseeding at 1:10 dilution. At all steps, media volumes were 0.2 mL/cm^2^.

After passaging, the cells were reseeded on fresh Matrigel-coated plates. Daily media changes were performed with mTeSR™1 medium, typically ± 3 h from the initial seeding time. After three to four rounds of clump passaging, pancreatic differentiation was initiated. Human islets were obtained from the Alberta Diabetes Institute Islet Core.

Human islets for research were provided by the Alberta Diabetes Institute IsletCore at the University of Alberta in Edmonton (www.bcell.org/adi-isletcore) with the assistance of the Human Organ Procurement and Exchange (HOPE) program, Trillium Gift of Life Network (TGLN), and other Canadian organ procurement organizations. Islet isolation was approved by the Human Research Ethics Board at the University of Alberta (Pro00013094) and by the University of McGill (22-02-054). All donors’ families gave informed consent for the use of pancreatic tissue in research.

### 1.2. Stem cell-derived pancreatic islet differentiation

The pancreatic differentiation protocol was carried out as previously described ^17–20^, with minor modifications. The H1 cells used for this process had a passage number between 30 and 40. Culture surfaces were pre-coated with growth factor-reduced Matrigel (Corning™, 35623) for 120 min. A seeding density of 130,000 cells/cm^2^ and 0.21 mL/cm^2^ mTeSR™1 medium was used. Media changes were performed daily from stage 1 to stage 6, and every other day during stage 7, typically ±1 h from the previous feeding time. At stage 4, day 4, aggregates were formed using the AggreWell™ 400 6-well plate (STEMCELL™ Technologies, 34425). Cells were harvested from the culture plate surface using TrypLE™ Select Enzyme (1X) (Thermo Fisher Scientific, 12605010). Three seeding densities were evaluated: 500, 1000, and 1500 cells per microwell, with later studies conducted at 1000 cells/microwell. At stage 5, day 7, the aggregates were transferred to ultralow attachment plates (Corning, CLS3471-24EA) and placed on a CellTron CO_2_ resistant shaker (Infors, 510925) at 100 rpm rotational speed, placed inside a humidified 5% CO_2_ incubator. Aggregates were encapsulated on day 7 of stage 6, followed by another transfer to ultralow attachment plates and placement on a Celltron orbital shaker at 100 rpm (for Objectives 1 and 2) or in a 100 mL PBS mini bioreactor at an agitation rate of 60 rpm ^20^. The bioreactors were seeded at a cell concentration of 1.5 ×10^5^ cells/mL. Every 48 h media changes were performed with stage 7 complete medium, typically ± 3 h from the initial seeding time. Figure 1 shows the schematic representation of the pancreatic differentiation protocol. The media formulation, growth factors, and small molecules used are detailed in Supplement Table 1.

**Figure 1.**
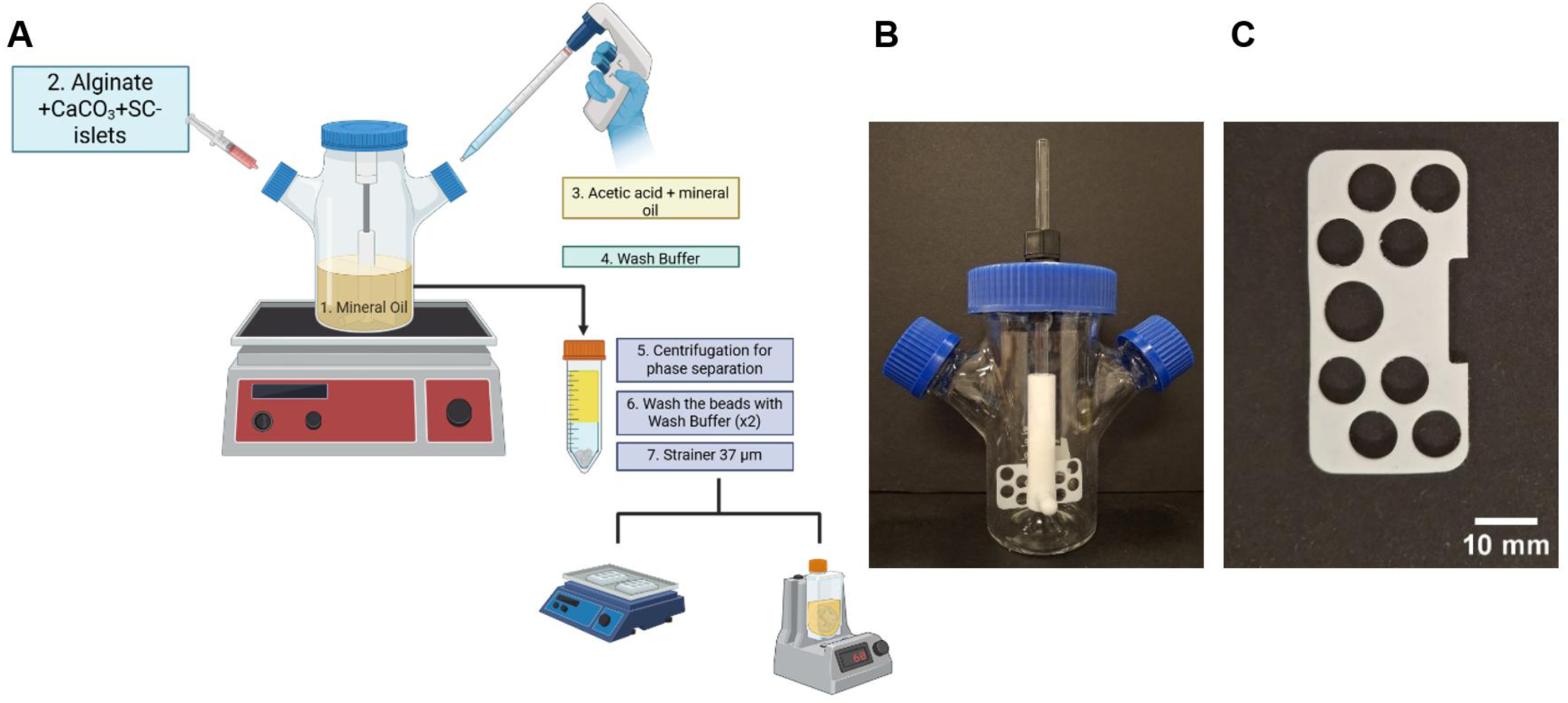
Emulsion-based encapsulation device and the microbead production process. **(A)** Schematic drawing of the SC-islets emulsion-based encapsulation process. Created with BioRender.com **(B)** Photograph of the emulsion-based encapsulation device (Micro-carrier Spinner Flask, BELLCO, 1965-00100). **(C)** Photograph of the impeller used in the emulsion-based encapsulation process.

### 1.3. Hydrogel preparation

A 50:50 mixture of the ultrapure sodium alginate PRONOVA^TM^ UP MVG (Novamatrix, Catolog #4200101) and PRONOVA^TM^ UP LVM (Novamatrix, Catolog # 42000001) was used to prepare 2.43%, 6.0% and 8.5% alginate stock solutions to respectively obtain 2%, 5% and 7% alginate beads (notwithstanding swelling). The alginate powder was dissolved in HEPES buffered saline solution consisting of 10 mM 4-(2-Hydroxyethyl) piperazine-1-ethanesulfonic acid, N-(2-Hydroxyethyl) piperazine-N′-(2-ethanesulfonic acid (HEPES; Thermo Fisher Scientific, BP310-500), and 170 mM NaCl (Sigma Aldrich, S9888) at pH 7.4, and then autoclaved at 121°C in a humid vapor cycle for 30 min (STERIS, AMSCO® Lab 110-250).

### 1.4. Emulsion-based encapsulation and internal gelation

The emulsion-based encapsulation and internal gelation method previously described ^15,16^ was followed with some modifications. Immature SC-islets (stage 6 day 7) were used in the encapsulation process.

An aqueous cell mixture was prepared as follows: 1.24 mL of alginate stock solution, 130 µL of 0.5 M CaCO_3_ solution in 10 mM HEPES buffer (Thermo Fisher Scientific, BP310-500), and 137 µL of SC-islets suspension (2.0×10^4^ cells/µL) in Stage 7 medium (Table 1). The mixture was gently mixed using a stainless-steel laboratory spatula.

Frist, 40 mL of light mineral oil (Thermo Fisher Scientific, O121-1) was agitated at 300 rpm for 2 min in a 100 mL microcarrier spinner flask (Bellco, 1965-00100) with a modified perforated impeller (Figure 2C). After this initial agitation, the stirring speed was reduced to 210 rpm. The aqueous cell mixture was then added dropwise to the spinner flask and emulsified for 12 min.

**Figure 2.**
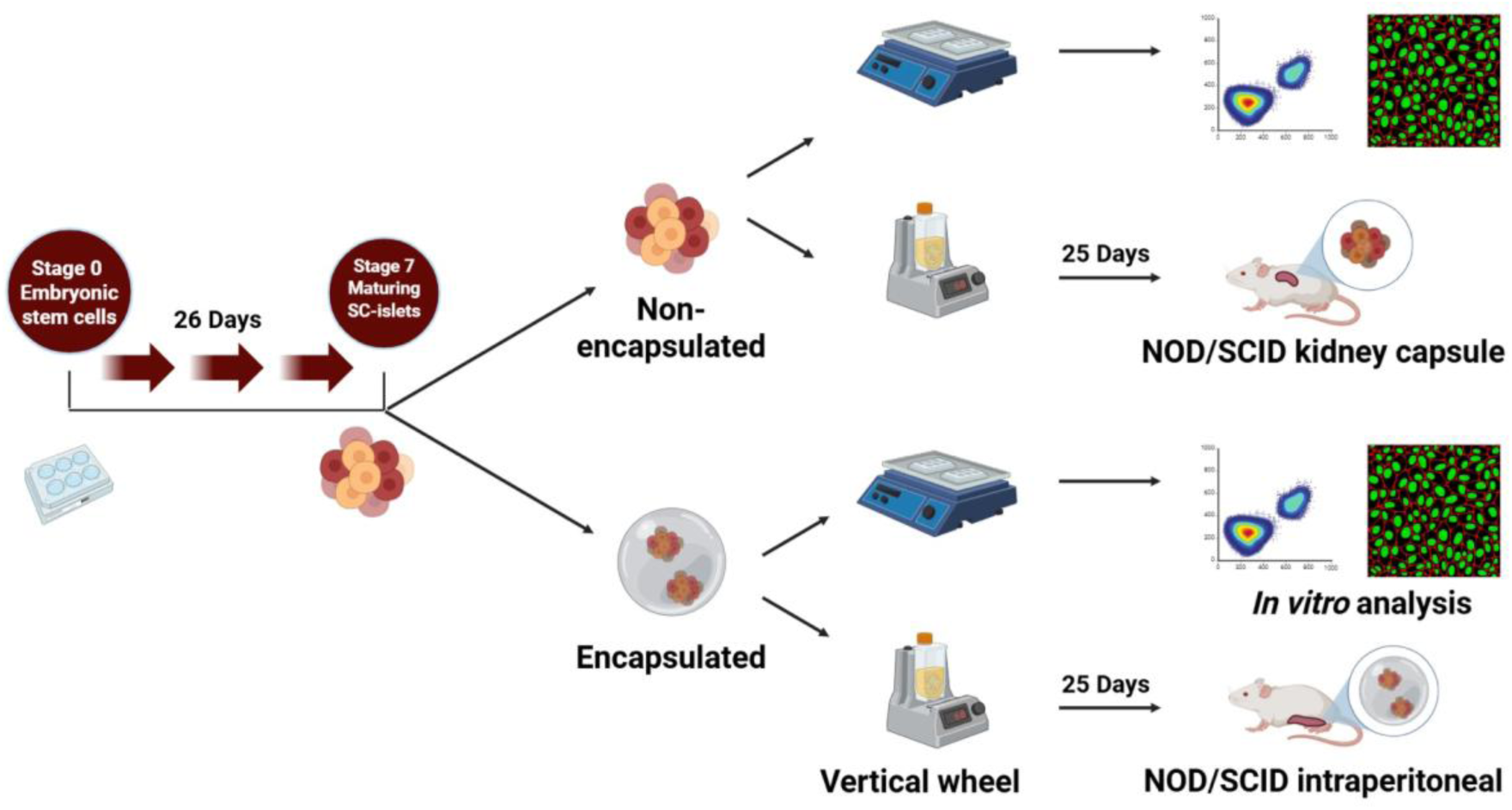
Schematic representation of the pancreatic differentiation protocol, encapsulation, culture and transplantation workflow. Created with BioRender.com

A limitation of the higher alginate concentration condition (7% w/v alginate) was that the spinning speed could not be increased beyond a certain threshold without compromising cell viability. Consequently, the maximum agitation rate tolerated by the cells resulted in a volume moment mean diameter (D4,3) of 1701 μm (SEM 76 μm). To enable meaningful comparisons between formulations, the agitation rate for each condition was adjusted to achieve a consistent D4,3 across all bead batches. The stirring speeds used for the 2%, 5%, and 7% alginate formulations were 240, 380, and 460 rpm, respectively, resulting in beads with a similar D4,3 value. The mixing conditions were optimized to maintain adequate cell viability while preserving the target bead diameter, particularly for the 7% alginate formulation.

Beads were stained with 1% toluidine blue (Sigma-Aldrich, T3260-5G) and subsequently imaged using a Samsung Galaxy A32 smartphone. The equivalent diameter based on the projected surface area in 2D images was measured using the Analyze Particles function in ImageJ^21^. The D4,3 values and standard errors of the mean (SEM) were calculated from the bead diameter distributions of each batch using Microsoft Excel.

Following the emulsification, 10 mL of acidified light mineral oil (9.64 µL glacial acetic acid dissolved through vortexing in 10 mL light mineral oil immediately prior to starting the process) was added to the spinner flask to trigger internal gelation. After 8 min, 40 mL of wash buffer (99.3 mM CaCl_2_, 75.0 mM NaCl, and 10.0 mM HEPES, pH 7.4) mixed with 10% stage 7 media was added to neutralize the pH. The oil, beads, and aqueous solution from the spinner flask were retrieved and collected in 50 mL Falcon tubes for centrifugation at 300 × g for 5 min. The oil was removed by aspiration through a Pasteur pipette with tip placed near the interface. Two further washes were performed. The beads were strained using a 37 µm reversible nylon mesh strainer (STEMCELL Technologies, 27250). The encapsulated SC-islets were further transferred to ultralow adhesion 6-well plates or to a 100 mL PBS mini bioreactor (1.5–2.0 × 10⁵ cells/mL). Figure 2 shows a schematic representation of the SC-islets emulsion-based encapsulation process.

### 1.5. Cell quantification

A degelling solution was prepared using 55 mM sodium citrate tribasic dihydrate (Sigma Aldrich, C8532), 90 mM sodium chloride (Sigma Aldrich, S5886), 10 mM HEPES (Thermo Fisher Scientific, BP310-500), and adjusted to pH 7.4. Immobilized SC-islets were recovered by incubating the alginate beads in 25 mL of degelling solution per gram of beads under agitation at 100 rpm on ice. After the incubation period, 25 mL of Stage 7 medium was added to dilute the degelling solution, and the suspension was centrifuged at 300 × g for 5 min. Harvested clusters were washed twice with DPBS without Ca^2+^ or Mg^2+^. To dissociate the aggregates into single cells, the clusters were incubated in TrypLE for 8–10 min at 37°C. The TrypLE was then diluted with Stage 7 medium, and the suspension was centrifuged again at 300 × g for 5 min ^22^. Following dissociation, the single cells were resuspended in fresh Stage 7 medium for cell counting. Both manual counting (using a Bright-Line™ Hemacytometer, Sigma Aldrich) and automatic counting (using a TC20 Automated Cell Counter, Bio-Rad) were performed to quantify cell concentration.

### 1.6. Flow cytometry

Cell monolayers or clusters were harvested and washed twice with DPBS (without Ca^2+^ or Mg^2^), then dissociated into single cells using TrypLE. Clusters were incubated at 37°C for 8-10 min and further broken apart through gentle pipetting up and down at least five times. After dissociation, single cells were resuspended in FACS Buffer (DPBS supplemented with 2% FBS ^22^). Next, the cells were incubated in the dark for 30 min with fixable viability dye (Life Technologies, L34963), washed twice with FACS Buffer, and fixed and permeabilized using Cytofix/Cytoperm (BD Biosciences, 554714) for 10 min.

The conjugated antibodies used are detailed in Table 4. Fixed cells were incubated with the conjugated antibodies for 30 min in the dark at 4°C, washed twice with Perm/Wash buffer (BD Biosciences, 554723), and then resuspended in FACS buffer for analysis. Flow cytometry was performed using BD Accuri^TM^ C6 Plus Flow Cytometer, and the data were analyzed using FlowJo v10 (BD Biosciences).

### 1.7. Dithizone staining

Dithizone powder (Sigma-Aldrich, 43820) was reconstituted in 99.7% Dimethyl sulfoxide (DMSO) (Fisher Scientific, BP231-100) (0.5mg/mL) and subsequently diluted in 4 mL DPBS (without Ca^2+^ or Mg^2^) to a final concentration of 0.125 mg/mL. The solution was filtered using 0.2 µm syringe filters (Fisher Scientific, 13100106) to eliminate any particulate matter. The aggregates were stained in the Dithizone solution for 1-2 min, followed by multiple rinses (∼10) with Ca^2+^ and Mg^2+^ free DPBS. Images were captured using a Zeiss Stemi 2000-C Stereo Microscope (455053) and Samsung Galaxy A32 smartphone.

### 1.8. Live/Dead staining

Encapsulated and non-encapsulated SC-islets were incubated for 20 min at 37°C using a live/dead staining solution containing 18.9 μg/mL propidium iodide (Fisher Scientific, P1304MP) and 1.1 μg/mL Calcein AM (Fisher Scientific, C3099). The clusters were then visualized, and images were acquired using IX81 Olympus Microscope with the FITC filter cube (Ex: 482/35 | Em: 536/40) and the Texas Red filter cube (Ex: 525/40 | Em: 585/40) ^22^. ImageJ software was utilized for image processing.

### 1.9. Static glucose-stimulated insulin secretion (GSIS)

To assess the SC-islets’ ability to secrete insulin in response to varying glucose concentrations, 20 encapsulated or non-encapsulated SC-islets, as well as cadaveric human islets, were hand-picked. The clusters were first incubated for 1 h in Krebs buffer supplemented with 2.8 mM glucose to equilibrate the system to basal glucose levels ^23^. The basal Krebs buffer (no glucose) was prepared as follows: 129 mM NaCl (Sigma Aldrich, S9888), 4.7 mM KCl (Sigma Aldrich, P3911), 2.5 mM CaCl_2_·2H_2_O (Sigma Aldrich, Catalog # 223506), 1.2 mM MgSO_4_ (Sigma Aldrich, 7487-88-9), 1.2 mM KH_2_PO_4_ (Sigma Aldrich, 7778-77-0), 5.0 mM NaHCO_3_ (Thermo Fisher Scientific, 144-55-8), 10 mM HEPES (Thermo Fisher Scientific, Catalog # BP310-500), 0.1% BSA (Sigma Aldrich, A3294), pH 7.4.

Subsequently, the clusters underwent a series of incubations: first in Krebs buffer supplemented with 2.8 mM glucose (low glucose incubation, 1 h), followed by Krebs buffer supplemented with 16 mM glucose (high glucose incubation, 1 h), then Krebs buffer with 2.8 mM glucose again (second low glucose incubation, 1 h), and finally Krebs buffer supplemented with 30 mM KCl (1 h). Two washes with Krebs buffer supplemented with 2.8 mM glucose were performed between the high glucose and second low glucose incubations. All samples and their respective pellets were stored at −20°C until analysis. DNA quantification was performed using Quant-iT™ PicoGreen™ dsDNA Assay Kits and dsDNA Reagents (Invitrogen™, P11496).

Human C-peptide levels were measured by ELISA (ALPCO human C-peptide kit, 80-CPTHU-E01.1) following the manufacturer’s instructions, as indication of SC-islet derived insulin secretion. Absorbance (450 nm) of all samples was measured using the Benchmark Plus Microplate Spectrophotometer (Bio-Rad).

### 1.10. Mechanical properties of the beads

To analyze the mechanical properties and viscoelasticity of different alginate bead formulations (2%, 5%, and 7%), the protocol described by Shin *et al.* ^24^ was followed. The MicroSquisher (CellScale), equipped with a parallel-plate compression configuration and a 559 μm diameter microbeam, was used for testing. Beads with the following size ranges were evaluated: 1100–1700 μm (2% alginate), 950–1100 μm (5% alginate), and 875–1200 μm (7% alginate). The beads were placed on a metal platform inside a chamber filled with Wash Buffer solution (see Section 4.4) at room temperature. To measure force as a function of displacement, the SquisherJoy software was utilized. Each bead underwent three consecutive cycles of up to 30% volume compression, with the following time settings: 30 seconds of loading, 10 seconds of holding, and 30 seconds of recovery. The compressive modulus (E) for each bead at 10%, 20%, and 30% compressive volume was calculated using MATLAB software and the Hertzian half-space contact model (refer to Equations 1, 2, 3, and 4) ^20,21^.

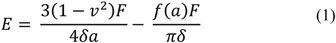

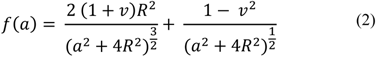

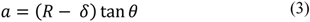

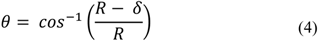

Where *R* is the radius of the bead, *v* is Poisson’s ratio, *δ* is the compressive displacement, *F* is the applied force during compression and *E* is the Young’s modulus.

### 1.11. Aggregate size measurement

To assess the aggregate size at various stages, brightfield images were captured using a VWR® Trinocular Inverted Microscope. Image analysis of the aggregates’ size was conducted using Fiji/ImageJ (NIH). First, edge detection was performed to enhance object boundaries, followed by conversion to a binary mask. Internal gaps within segmented objects were filled using the Fill Holes function. Adjacent objects were then separated using the Watershed algorithm. Quantitative measurements, including area, mean intensity, minimum intensity, perimeter, shape descriptors, and Feret’s diameter, were extracted for each object. Particle analysis was subsequently performed using a minimum size threshold of 600 µm², and only particles meeting this criterion were included in the analysis. Object outlines were generated for visualization, and summary statistics were exported for further analysis.

### 1.12. Transplantation studies

Murine transplantation experiments were performed at McGill University following a protocol approved by the McGill Facility Animal Care Committee (Protocol AUP# 8116). Six- to nine-week-old male immunocompromised mice (NOD.Cg-Prkd<scid>/J; The Jackson Laboratory) weighing 24 to 27 g were anesthetized with inhalable isoflurane. Stage 7 Day 25 SC-islets from the same batch were transplanted at doses equivalent to 2 million cells per mouse. Non-encapsulated SC-islets were transplanted into the kidney capsule while encapsulated SC-islets suspended in 0.3 mL serum-free Dulbecco’s Modified Eagle Medium (DMEM, ThermoFisher; 11054020) were injected into the intraperitoneal space using a sterile 1 mL pipette with cut tip.

Functional assessment included measurement of human C-peptide levels and meal challenge. For routine monitoring, mice were fasted for 4h every two weeks before bloo collection. At the designated meal challenge time point, mice were fasted for 16 h and fed during 45 min. Blood samples (80 µL) were collected by saphenous vein blood collection into microvettes (Kent Scientific; MCVT100-LIHEP). Whole blood was centrifuged for 9 minutes at 4000 x g at 4°C, and serum supernatant was collected and stored at −80°C for human C-peptide quantification by ELISA. Body weight was measured throughout the study. For the kidney capsule group, three transplantations were attempted, but in one case the free islets were not successfully delivered into the capsule at the time of transplantation, so only two mice were included in the final analysis for this group.

### 1.13. Immunohistochemistry

Encapsulated islets or excised kidneys were washed with HEPES solution and then fixed in Bouin’s solution (8.8% formaldehyde and 5% glacial acetic acid). After 15 min of fixing on ice at 50 rpm on a rotary shaker, the beads were washed with HEPES solution before paraffin embedding and sectioning at4 µm. Kidneys were fixed overnight in 4% paraformaldehyde, embedded in paraffin and sectioned at 4 µm. Paraffin sections were rehydrated by sequential 5 min incubations in xylene (three times), 10 min in 100% ethanol (twice), 95% ethanol, and then 70% ethanol. The sections were washed in PBS and microwaved (800W) for 17 min in 10 mM citrate buffer with 0.05% Tween 20 at pH 6.0. Samples were blocked with 0.1% Triton X-100 (Sigma; T8787) and 5% bovine serum albumin (BSA, Sigma; A7906) in PBS (staining solution) for 1h at room temperature (RT), incubated with primary antibodies (mouse anti-glucagon from Sigma, 1:200 dilution; G2654 and guinea pig anti-C-peptide from Abcam 1:100 dilution; ab30477) diluted in staining solution overnight at 4°C, washed for 10 min in rinsing solution (0.1% Triton X-100, 0.1% BSA in PBS), incubated with appropriate Alexa Fluor-488 or −555 secondary antibodies (Thermo Fisher; a11073 and a21424) diluted 1:500 in staining solution 1h at RT, washed for 10 min in rinsing solution, mounted with Vectashield with 4′,6-diamidino-2-phenylindole (DAPI,Cedarlane; VECTH1200) and covered with a coverslip. Images were taken with an LSM 710 Confocal Microscope (Zeiss) equipped with the Zeiss FS 49 (DAPI), Zeiss FS 38 (eGFP), and Zeiss FS 45 (mCherry) fluorescence filter sets.

### 1.14. Bulk RNA-sequencing analysis

SC-islets were collected and digested in RLT lysis buffer from the RNeasy Kit (Qiagen, 74104) supplemented with beta-mercaptoethanol (Sigma) and stored in −80 °C. RNA extraction was performed using the above kit. RNA-seq was performed by the Molecular Biology and Functional Genomics Platform of the Montreal Clinical Research Institute. RNA libraries were prepared from 1000 ng of total RNA. The mRNA was enriched using the NEBNext® Poly(A) mRNA Magnetic Isolation Module (New England Biolabs) and libraries were prepared with RNA Hyperprep Kit (KAPA). Library size distribution was assessed on a 2100 bioanalyzer (Agilent Technologies) and libraries were quantified by qPCR (QuantStudio 5, Life Technologies). Equimolar libraries were sequenced in paired-end reads (PE100), on a Novaseq X system (Illumina), with a coverage of 50M fragments per library. To generate the count matrix, raw FASTQ files were processed using the standard nf-co.re RNA-seq pipeline (https://nf-co.re/rnaseq/3.15.0) with the star salmon workflow to align to the human genome build (hg38). Analysis was performed in RStudio (v4.4.0). Differential gene expression analysis was performed using the DESeq2 R package (1.20.0) with an adjusted p-value <0.05 considered significant. Gene ontology term analysis was performed using EnrichR with default settings, with all annotated genes in the KEGG_2021_Human^25^.

### 1.15. Statistical analysis

Aside from RNAseq studies, statistical analyses were performed using GraphPad Prism 10. Data normality was assessed using the Shapiro-Wilk test, and P-values < 0.05 were considered statistically significant. For parametric data, one-way ANOVA with Tukey’s multiple comparisons test was used. Non-parametric data were analyzed using the Brown-Forsythe ANOVA test with Kruskal-Wallis with Dunn’s multiple comparison test. Graphs were generated using GraphPad Prism and Microsoft Excel. Data are presented as the mean ± standard error of the mean (SEM) of independent experimental replicates.

## 2. Results

### 2.1. Effect of aggregate size and time on SC-islet agglomeration and yields

To assess the effect of initial cell aggregate size on subsequent agglomeration in suspension, 500, 1000 or 1500 stage 4 cells were seeded per microwell at the start of stage 5. After 5 days of microwell culture to stabilize size, the aggregates were transferred into suspension culture for 17 days (10 days stage 6 + 7 days stage 7). As expected, the volume estimated from the cross-sectional area of aggregates increased nearly proportionally to the number of cells seeded per microwell (Figure 3B left). After 7 days of stage 6 suspension culture, aggregate diameters increased to 124 µm (SD 2.4) and remained constant up to Stage 7 day 10 (Figure3 B), irrespective of the initial number of Stage 4 cells seeded per microwell. Substantial cell loss was observed both during microwell aggregation and subsequent suspension culture, resulting in yields of 36% (SEM 2%) at Stage 5 day 5 and 14% (SEM 1%) at Stage 7-day 10 relative to the Stage 4 Day 3 seeding density (Figure 3C). Stage 7 day 10 cultures yielded 60% (SEM 1%) NKX6.1+, 75% (SEM 2%) NeuroD1+, 19% (SEM 1%) C-peptide+ cells, 19% (SEM 1%) glucagon, with no significant differences between initial aggregate sizes at stage 4 (Figure 3D). The midpoint condition (1000 cells/aggregate) was applied in subsequent experiments.

**Figure 3.**
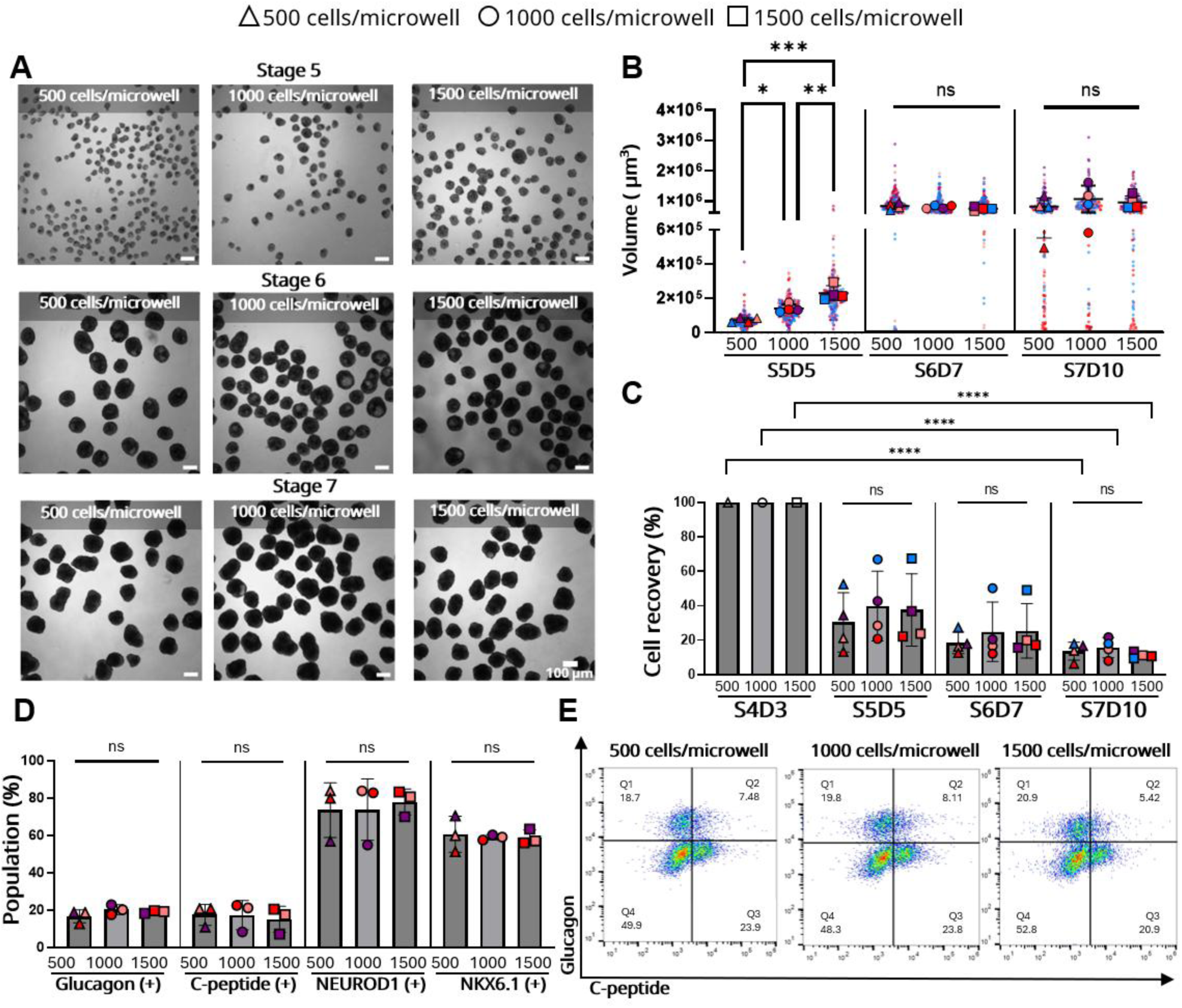
Impact of seeding density (500, 1000, and 1500 cells/microwell) during pancreatic progenitor aggregate formation on aggregate size, cell recovery and expression of pancreatic markers. **(A)** Morphological images and aggregate sizes at stage 5, 6, and 7 for each condition. **(B)** Graph showing size distribution across stages for each condition. Data is presented as the mean and standard deviation (N=4, n=60). *p < 0.05, **p < 0.01, ***p < 0.001 by one-way ANOVA with Tukey’s multiple comparisons test (Stages 5 and 6) and ns = not significant by Kruskal-Wallis with Dunn’s multiple comparison test (stage 7). Ns = not significant by one-way ANOVA with Tukey’s multiple comparisons test. S5D5 = stage 5 day 5, S6D7 = stage 6 day 7, S7D10 = stage 7 day 10. **(C)** Percentage of total cell recovery normalized by seeding density during aggregate formation (stage 4) at each stage (N=4). ****p < 0.0001 by two-way ANOVA with Tukey’s multiple comparisons test. **(D)** Flow cytometry analysis of pancreatic marker expression at stage 7 (day 10) of maturation. Statistical non significance was determined by one-way ANOVA with Tukey’s multiple comparisons test. **(E)** Representative flow cytometry plots of glucagon versus C-peptide.

### 2.2. Encapsulation to Minimize Cluster Aggregation and Reduce Cell Loss

We hypothesized that cellular agglomeration and cell losses from hydrodynamic stress could be reduced through microencapsulation. As the effect of different alginate concentrations on SC-islet differentiation was unknown, we generated 2%, 5% and 7% alginate beads. The midpoint, 5% alginate with equal mass concentration of LVM and MVG alginate, was previously identified as promising in protecting beta cells from allogeneic rejection in mice^26^. As higher alginate concentrations increase fluid viscosity, the mixing speed during emulsification was adjusted to achieve similar D_43_ values between conditions (Figure 4A). Given the polydisperse size distribution of alginate beads produced by stirred emulsification, D_43_ values reflect the relative frequency of cells in beads of different diameters. The mechanical properties of representative hand-picked ∼800 μm diameter beads were characterized using the Hertzian contact model to determine the compressive Young’s modulus (E) at 10%, 20%, and 30% deformation. As expected, capsule stiffness increased with alginate concentration, with higher Young’s modulus values measured for 5% and 7% alginate compared to 2% (Figure 4B).

**Figure 4.**
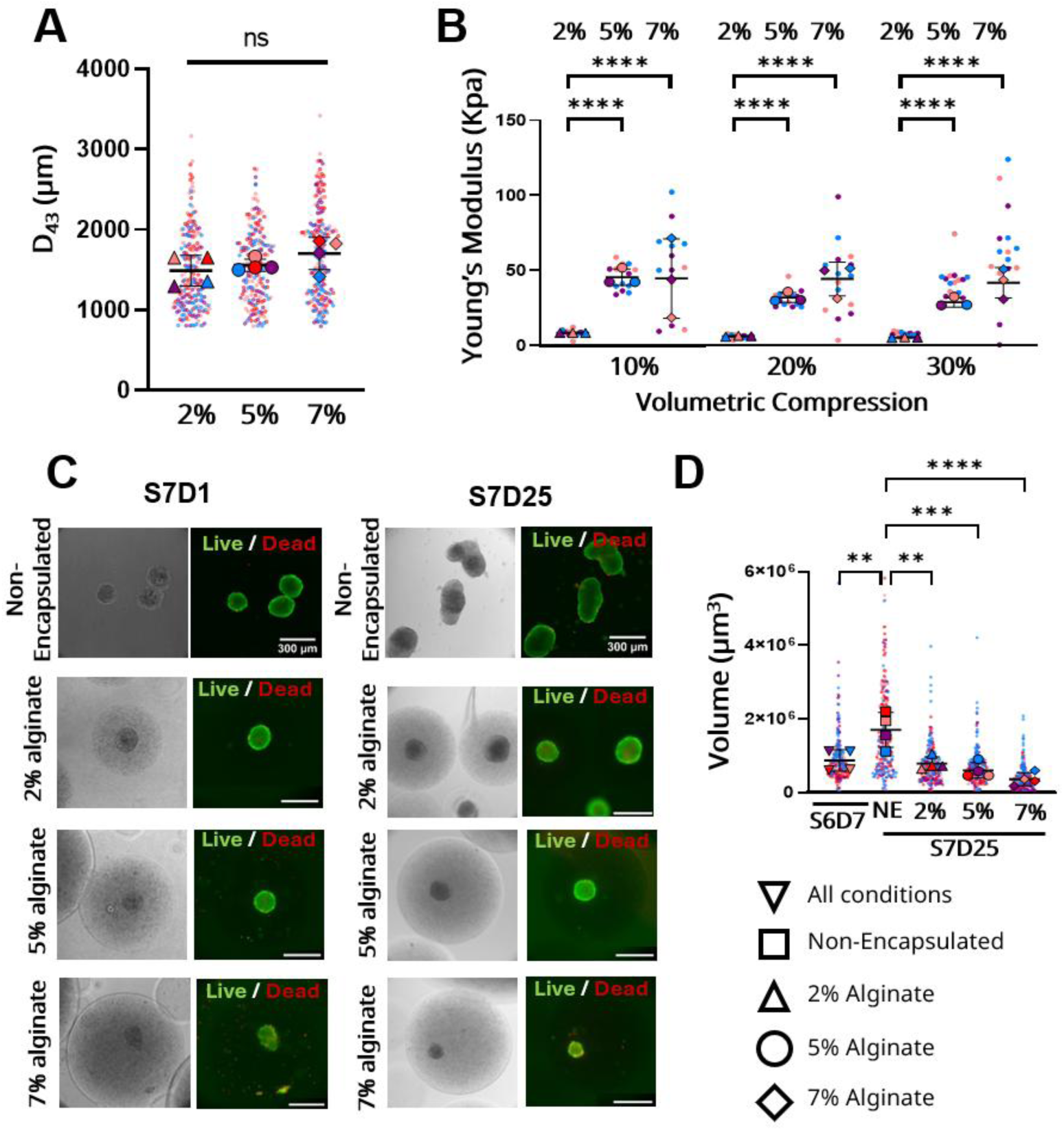
Emulsion-based encapsulation. **(A)** Volume moment mean diameter (D_43_) of the beads produced with 2, 5, and 7% alginate. N=4, n=50. **(B)** Compressive modulus of beads at 10%, 20%, and 30% compressive volume for different alginate concentrations (N=3, n=15). **(C)** Fluorescence images of Live/Dead staining of stage 7 Day 1 immature SC-islets, 24 hours post-encapsulation using Calcein-AM (Green) and Propidium Iodide (Red) and Live/Dead staining of Stage 7 Day 25 maturing SC-islets, 25 days post-encapsulation. **(D)** Cluster size distribution before encapsulation (S6D7), and 25 days after encapsulation (S7D25) (Aggregates recovered from degelled beads). Data is presented as the mean and standard deviation (N=4, n=60). For all data shown, between-group comparisons were performed via one-way ANOVA with Tukey’s multiple comparisons test, with the exception of panel B at 10% and 20% compressive volume which used (10% and 20% compressive volume) and Brown-Forsythe ANOVA test and Games-Howell’s multiple comparisons test (30% compressive volume). **p < 0.01, ***p < 0.001, ****p < 0.0001 by one-way ANOVA with Tukey’s multiple comparisons test. S6D7 = stage 6 day 7, S7D25 = stage 7 day 25. S = Suspension (non-encapsulated).

SC-islets were encapsulated at the end of Stage 6 prior to maturation. Earlier encapsulation attempts resulted in significant cell loss, supporting the selection of this stage for encapsulation (Figure S2). SC-islet viability, morphology, and size were assessed following encapsulation and after 25 days of immobilized culture. Although some cell loss occurred during the encapsulation procedure itself, post-encapsulation viability remained high, and cluster morphology was preserved across all conditions.

At 7% alginate, SC-islets appeared qualitatively smaller immediately following encapsulation, likely due to increased shear associated with the higher mixing intensity required during bead production, although viable clusters were still observed (Figure 4C). Aggregate size showed little change over time in encapsulated conditions, whereas non-encapsulated controls exhibited a trend toward increased aggregate size consistent with agglomeration. Collectively, these findings are consistent with encapsulation limiting aggregate fusion during maturation and promoting greater size uniformity over time (Figure 4D).

### 3.3 Effect of Alginate Stiffness on Pancreatic Differentiation

To evaluate if the encapsulation microenvironment had an impact on the maturation of SC-islets, the expression of several pancreatic markers was assessed: glucagon (associated with alpha-cells), C-peptide (associated with beta-cells), NEUROD1, NKX6.1, and PDX1 (key transcription factors involve in the development and functionality of different endocrine cells which comprise the islets of Langerhans^27^). Figure 5A shows that SC-islets encapsulated in 7% alginate contained a significantly higher fraction of glucagon-positive alpha-cells than those encapsulated in 2% alginate or non-encapsulated SC-islets. Conversely, the differentiation of beta-cells, as indicated by C-peptide expression, and the expression of NKX6.1 were favored in non-encapsulated environments, which had lower stiffness, compared to the high-stiffness environment (7% alginate). The expression of other pancreatic markers, NEUROD1 and PDX1, had no significant differences across all conditions.

**Figure 5.**
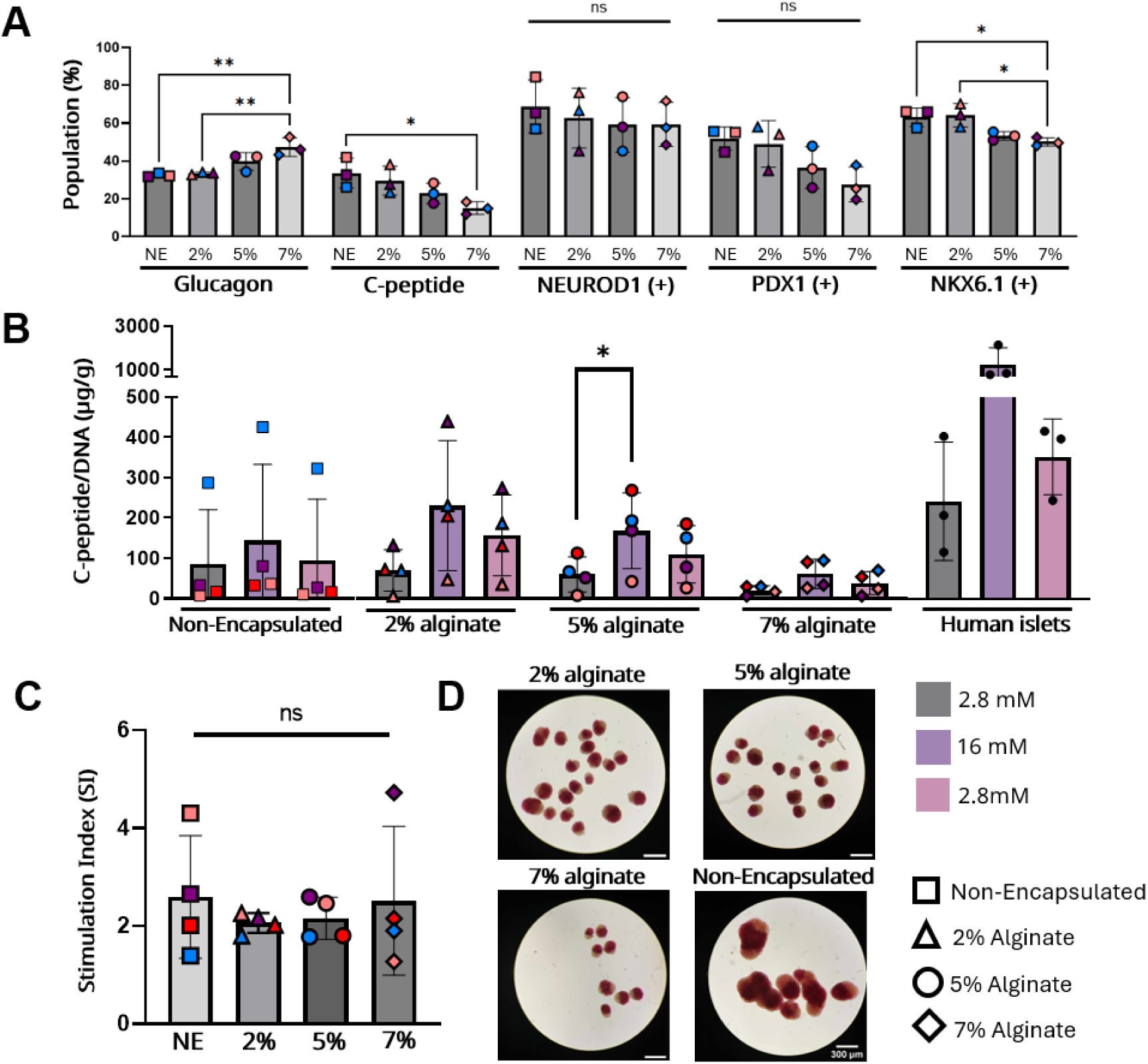
Effect on alginate stiffness on pancreatic differentiation **(A)** Flow cytometry analysis of pancreatic marker expression at stage 7 (day 25) of maturation. *p < 0.5, **p < 0.01, ns = not significant by one-way Anova and Turkey’s multiple comparisons test (for Glucagon, C-peptide, NKX6.1, PDX1), and Kruskal-Wallis for NEUROD1 with Dunn’s multiple comparison test (N=3). **(B)** Static Glucose-Stimulated Insulin Secretion (GSIS) of mature SC-islets after 3 weeks of culture in alginate beads with varying alginate concentrations (cells encapsulated at stage 6 day 7) (N=4) and human islets (N=3). *p < 0.5 by two-way ANOVA with Tukey’s multiple comparisons test. **(C)** Stimulation Index of maturing SC-Islets after 3 weeks of encapsulation. Ns = not significance by one-way ANOVA with Tukey’s multiple comparisons test (N=4). **(D)** Dithizone staining of maturing SC-islets, encapsulated and non-encapsulated after 3 weeks culture at different alginate concentrations (aggregates recovered from degelled beads).

The functionality of the maturing SC-islets was evaluated *in vitro* through a static glucose-stimulated insulin secretion assay (Figure 5B). As expected, SC-islets showed increased insulin secretion in response to glucose, with those immobilized in softer environments exhibiting higher levels of insulin secretion. Compared to human islets, the SC-islets displayed a lower insulin secretion response, indicating that while the necessary machinery for glucose-responsive insulin secretion is present, these stem cell-derived clusters remain relatively immature compared to fully mature human islets. Among the SC-islet groups, clusters immobilized in 7% alginate beads exhibited lower absolute insulin secretion than the other conditions. This diminished response correlates with the lower fraction of beta cells present in these beads after 25 days (Figure 5A).

Notably, despite differences in the magnitude of insulin secretion, all SC-islet groups exhibited appropriate glucose-responsive behavior, with insulin secretion increasing in response to high glucose (16 mM) and returning to basal levels upon re-exposure to low glucose (2 mM). The ability of the SC-islets to reduce insulin secretion upon return to low glucose further demonstrates preserved glucose-sensing dynamics. Consistent with these observations, the insulin stimulation index, calculated as the ratio of insulin secreted in response to high glucose relative to the average insulin secretion during the two low-glucose exposures, did not differ significantly between conditions (Figure 5C). These findings suggest that encapsulation did not impair the functional glucose responsiveness of the SC-islet clusters.

SC-islets were further characterized by staining with DTZ, a dye which interacts with zinc present in insulin granules (Figure 5D). Consistent with previous observations, SC-islets maintained in suspension displayed the characteristic agglomeration phenotype. Interestingly, regions of high-intensity DTZ staining appeared spatially segregated from areas with lower DTZ intensity, a pattern previously reported in endocrine cell ‘budding’ models and suggestive of self-organization of cellular subtypes within the clusters^28^.

### 3.3 Scale-up of encapsulated SC-islet cultured in 100 mL vertical wheel bioreactors

To assess the translational potential of this microencapsulation strategy, we next evaluated whether encapsulation could facilitate SC-islet scale-up in vertical wheel bioreactor cultures by protecting pancreatic aggregates from shear-induced damage and preventing aggregate agglomeration during suspension culture. Given the potential for direct transplantation following culture, encapsulation was performed using 5% alginate, the formulation selected for subsequent studies.

Both encapsulated and non-encapsulated clusters were inoculated into 100 mL PBS Mini bioreactors (Figure 6A and 6B) at a cell concentration of 1.5–2.0 × 10⁵ cells/mL, and agitated at 60 rpm (following a previous published study ^20^). Viability was evaluated 24 h and 25 days post-encapsulation using Live/Dead staining with calcein-AM and propidium iodide (Figure 6C and 6D). Results indicated that cell viability remained high both before and after encapsulation in 5% alginate. Similarly, non-encapsulated clusters maintained high viability. As expected, a noticeable increase in the size of non-encapsulated clusters was observed, from an average volume of 2.3 × 10^6^ µm^3^ (SEM 1.9 × 10^5^ µm^3^) before encapsulation to an average volume of 1.6 × 10^7^ µm^3^ (SEM 9.1 × 10^5^ µm^3^), contrary to immobilized aggregates, average volume of 2.2 × 10^6^ µm^3^ (SEM 1.0 × 10^5^ µm^3^) (Figure 6E). Encapsulation helped maintain a more uniform cluster size over time, by preventing aggregate fusion. Additionally, non-encapsulated clusters exhibited a lower recovery rate, 60% (SEM 9%), compared to encapsulated clusters, 91% (SEM 2%), likely due to increased exposure to shear stress, leading to higher cell loss (Figure 6F).

**Figure 6.**
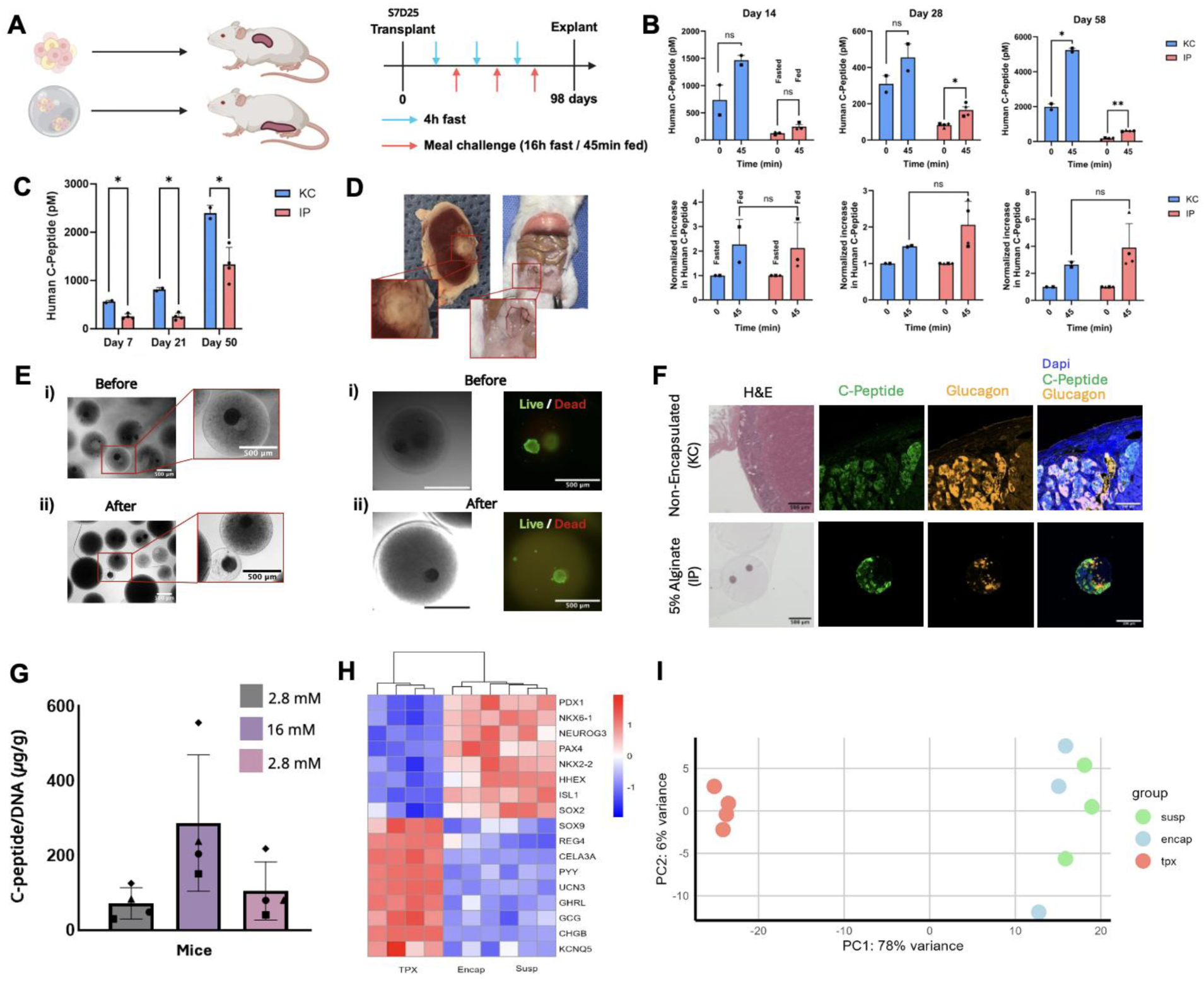
Transplantation of stem cell-derived islets (SC-islets) in non-diabetic mice using alginate microbeads. **(A)** Schematic summarizing *in-vivo* study. **(B)** Meal challenge (16 hours fast, 45 minutes fed) measurement at 16, 28, 58 days post-transplantation. *p < 0.05, **p < 0.01 by 2-way paired t-test comparing 0- and 45-minutes time points. **(C)** Serum human C-Peptide measurement after 4 hours fast at 7, 21, 50 days post transplantation. **(D)** Images of explanted grafts in kidney capsule (KC) and microbeads in intraperitoneal. **(E)** Encapsulated SC-islets in 5% alginate microbeads before transplantation **(i)** and after 98 days in vivo **(ii)** with live/dead staining using Calcein-AM (green) and Propidium Iodide (red). **(F)** Representative histology and immunofluorescence images of sectioned graft explanted from KC (non-encapsulated) and IP (5% alginate). **(G)** Static GSIS of encapsulated SC-islets after 98 days *in vivo*. **(H)** Principal component analysis (PCA) of the SC-islets cultured in suspension (non-encapsulated pre-transplant, S7D25), in encapsulation (encapsulated pre-transplant, S7D25), or transplanted *in vivo* (IP explant, S7D123). **(I)** Heatmap of selected differentially expressed genes (padj<0.05, log2FC>0.5) between the three conditions.

The maturation of stage 7 day 25 SC-islets was assessed by analyzing the expression of key pancreatic markers (C-peptide and glucagon), their functionality (static glucose-stimulated insulin secretion), and insulin storage (dithizone staining). Flow cytometry data revealed that C-peptide and glucagon expression remained consistent across both conditions (Figure 6G).

Both encapsulated and non-encapsulated SC-islets remained glucose responsive after 25 days of differentiation and maturation in 100 mL PBS mini-bioreactors. Although encapsulated SC-islets exhibited a trend toward lower insulin secretion (Figure 6H), one-way ANOVA detected no statistically significant differences in insulin secretion among culture groups under either low- or high-glucose conditions. It is important to note that these results were standardized based on DNA content. In the encapsulated condition, the DNA from dead cells remained trapped in the beads, while in suspension, the DNA from dead cells was removed during medium exchanges. This may have led to a less accurate DNA measurement, which could explain the observed differences.

Using bulk RNA-sequencing we look at differentially regulated genes between SC-islets cultured in suspension or encapsulated in the beads. Gene Ontology analysis (Figure 6J) showed that protein absorption and digestion as well as extracellular matrix (ECM)-receptor interaction are different between the two groups. Analysis of specific genes related to these gene ontology terms confirmed increased expression of extracellular matrix genes (COL2A1, LAMA3) and pancreatic enzymes (CPA1, CTRB2) in suspension culture, whereas matrix metalloproteinases (MMP) were elevated in encapsulated SC-islets (Figure 6K).

### 3.4 Long-term Survival and Function of Transplanted Encapsulated SC-islets

To assess the long-term viability and in vivo function of SC-islets obtained in bioreactors, Stage 7 SC-islets were transplanted intraperitoneally into immunodeficient mice following alginate encapsulation (Figure 7A). As a positive reference control, a small cohort of non-encapsulated SC-islets (n = 2 mice) was transplanted under the kidney capsule, the most widely used transplantation site for assessing islet graft function because of its highly vascularized and well-defined microenvironment. Since the kidney capsule cannot accommodate alginate microbeads due to its limited physical space, this control group was included to confirm the functionality of non-encapsulated SC-islets and provide a reference to the existing literature, rather than for a fully powered comparison between transplantation sites.

Encapsulated SC-islets transplanted intraperitoneally exhibited robust glucose-responsive function throughout the study. Human C-peptide levels increased after refeeding at multiple time points, demonstrating preserved glucose-stimulated insulin secretion in vivo (Figure 7B, top panels). Importantly, the normalized increase in C-peptide relative to fasting remained comparable to kidney capsule transplantation (Figure 7B, bottom panels), indicating that encapsulation did not impair glucose responsiveness following long-term transplantation.

Longitudinal monitoring further demonstrated sustained graft function, with detectable human C-peptide secretion as early as day 7 after transplantation that persisted throughout the 98-day study (Figure 7C). As expected, the kidney capsule reference animals exhibited higher absolute C-peptide concentrations, consistent with the superior vascularization of this transplantation site. However, because the kidney capsule cohort comprised only two animals, these observations are presented descriptively rather than as a formal comparison. The mean kidney capsule values fell outside the 95% confidence interval of the intraperitoneal group at each evaluated time point, suggesting greater secretory output in the kidney capsule. These differences should nevertheless be interpreted cautiously and warrant confirmation in larger studies specifically designed to compare transplantation sites^29^.

At day 98 post-transplantation, grafts were explanted for analysis (Figure 7D). Gross morphology showed no visible fibrotic overgrowth around the retrieved microbeads (Figure 7E ii), and Live/Dead staining confirmed that most of the encapsulated aggregates remained viable (Figure 7E, right panel). Immunofluorescence analysis of the explanted grafts confirmed preservation of endocrine differentiation within the retrieved aggregates, demonstrating maintenance of islet lineage identity after prolonged in vivo implantation. *Ex vivo* GSIS assays further demonstrated preserved glucose responsiveness of the explanted SC-islets (Figure 7G), indicated that endocrine identity was accompanied by maintained functional competence.

To assess molecular maturation status, transcriptomic profiling was performed on explanted SC-islets. Heatmap analysis (Figure 7H) demonstrated upregulation of mature β-cell markers alongside downregulation of progenitor-associated genes, consistent with continued maturation in vivo. Principal component analysis (Figure 7I) showed tight clustering of transplanted encapsulated samples along PC1 (78% variance), indicating strong reproducibility after transplantation. Explanted encapsulated SC-islets formed a distinct cluster separated from both non-encapsulated and encapsulated in vitro cultures, demonstrating convergence toward a stable and coherent *in vivo* transcriptional state.

These findings demonstrate that alginate encapsulation supports long-term viability and nutrient-responsive function of SC-islets *in vivo*, with no apparent fibrosis. Although vascularization in the intraperitoneal space may limit peak C-peptide levels, encapsulated SC-islets remained functional for over three months, highlighting their potential for clinically relevant transplantation strategies.

## 3. Discussion

This study is the first to demonstrate highly scalable one-pot emulsion microencapsulation of SC-islets in high-concentration alginate beads followed by extended *in vitro* culture and *in vivo* graft survival. Building on our earlier adaptation of alginate emulsification and internal gelation to mammalian cell including beta cell line immobilization^15,16^, here we show that this strategy can be applied to immobilize cell clusters such as SC-islets while maintaining viability, endocrine identity, and function. Stirred emulsification-based encapsulation offers several key features as compared to other islet encapsulation method. The process does not require specialized equipment, is not prone to nozzle or channel obstruction by cell clusters or debris, can readily be scaled up in closed systems, and is compatible with highly viscous solutions (up to 10 Pa·s^15^). Existing nozzle or microchannel-based systems often create beads with ripples or teardrop shapes through limited flight time or mechanical phenomena when contacting external gelation solutions^13,30^. Emulsification and internal gelation typically yields highly spherical beads with smooth surfaces because interfacial tension minimizes the surface area during droplet formation. This may present some advantages since bead imperfections and roughness can trigger foreign body responses^31,32^. These features enable islet encapsulation in high-concentration alginate beads with low permeability to antibodies, avoiding additional cationic polymer coatings applied to reduce the permeability of nozzle-generated hydrogel beads^16^.

SC-islets encapsulated in alginate concentrations up to 7% remained viable, retained key pancreatic markers, and responded appropriately to glucose stimulation. Previous studies reported that alginate encapsulation can enhance pancreatic differentiation, including increased PDX1 and NKX6.1 expression through integrin signaling, with outcomes dependent on the differentiation stage at encapsulation^33,34^. However, Legøy et al. used a substantially lower alginate concentration (1.8%) than those investigated here (2-5–7%), which may partly explain why a similar enhancement in PDX1 and NKX6.1 expression was not observed in our study. Interestingly, alginate concentration was associated with differences in SC-islet phenotype: higher alginate concentrations resulted in increased glucagon-positive populations and reduced frequencies of C-peptide- and NKX6.1-positive cells. Although these observations correlated with increased bead stiffness, multiple microenvironmental properties change simultaneously with alginate concentration, including matrix density, charge distribution, and mass-transfer characteristics. Differences in alginate concentration and encapsulation systems may therefore contribute to differences in endocrine marker expression between studies. Nevertheless, the reduced NKX6.1 expression and increased glucagon-positive fraction observed at higher alginate concentrations are consistent with developmental studies showing that alpha-cell precursors can arise from NKX6.1-negative progenitors and follow distinct lineage trajectories from beta-cells^35,36^. Additional transcriptomic studies will be required to determine whether these effects reflect altered endocrine fate decisions, selective survival of specific cell types, or differences in maturation kinetics.

A major finding of this study is that encapsulation prevented SC-islet agglomeration during extended suspension culture. Aggregate fusion observed in this work as well as prior studies with SC-islets and organoids cultured in suspension more broadly represents a significant challenge during organoid scale-up in suspension bioreactors^37–40^. Increasing cluster size exacerbates oxygen and other biochemical gradients which can adversely impact cell survival or differentiation outcomes^41,42^. Encapsulation stabilized the aggregate size distribution throughout maturation as compared to non-encapsulated controls which exhibited substantial enlargement through aggregate fusion visible through merged clusters (Figure 4C). Oxygen availability is known to play a critical role during pancreatic development and endocrine differentiation^43–46^, suggesting that the maintenance of smaller aggregate sizes may help preserve a more homogeneous culture environment. RNA sequencing further suggested differences in extracellular matrix remodeling and cell–matrix interactions between encapsulated and non-encapsulated cultures, indicating that the capsule microenvironment may actively influence cellular behaviour in addition to physically compartmentalizing aggregates. Encapsulation also markedly improved cell recovery over extended (up to 25 days) suspension culture both in agitated 6-well plates and vertical wheel systems. These findings suggest that alginate immobilization not only prevents aggregate fusion but may also provide protection against hydrodynamic stress during scale up.

An important advantage of emulsion-based encapsulation is the potential for integration with scalable and extended culture of SC-islets and likely other organoids. Cell immobilization facilitates cell retention and handling, for example by reducing settling times and preventing SC-islet agglomeration during frequent media exchange. The current microchannel emulsification and internal generation setup generates up to 10 mL total volume of alginate beads in less than 1h total processing time –with 10-20 mL beads theoretically sufficient to accommodate therapeutic doses for one recipient. Unlike nozzle or microchannel-based systems, scale-up of stirred emulsification entails a relatively simple increase in vessel diameter (proportional to the square root of the volume increase), rather than a linear increase in microchannel numbers or process duration. Encapsulated SC-islets were successfully matured in 100 mL PBS mini bioreactors while maintaining viability, pancreatic marker expression, and glucose responsiveness. Recent studies have demonstrated scalable production of non-encapsulated SC-islets in suspension and Vertical Wheel bioreactors while maintaining pancreatic identity and function^47,48^. Our results extend these approaches to encapsulated SC-islets, while additionally preventing aggregate fusion and improving cell recovery during extended culture. This is particularly relevant because future commercial production of SC-islets will likely rely on bioreactor technologies rather than static culture systems^23,49,50^.

Beyond its bioprocessing advantages, encapsulation has the potential to streamline the transition from SC-islet manufacturing to transplantation. The alginate formulation used in this study was selected based on previous reports demonstrating low fibrotic overgrowth and poor permeability to antibodies^16^, making it attractive for future immunoisolation applications. Previous work demonstrated long-term glycemic control following transplantation of alginate-encapsulated stem cell-derived beta cells in immunocompetent mice^51^. In the present study, encapsulation was instead integrated with scalable SC-islet processing and extended bioreactor culture prior to transplantation. Encapsulated SC-islets transplanted into immunodeficient mice secreted human C-peptide as early as seven days after transplantation, remained meal responsive, and continued to mature *in vivo*. To our knowledge, this is the first study demonstrating that SC-islets encapsulated in high-concentration alginate beads produced through a highly scalable one-pot emulsion process remain functional, glucose responsive, and capable of further maturation following transplantation. Although C-peptide levels remained lower than those observed under the kidney capsule, the intraperitoneal grafts demonstrated sustained viability and endocrine function throughout the 3-month transplantation period, supporting the feasibility of directly transplanting emulsion-encapsulated SC-islets following bioreactor culture.

Building on these promising bioreactor and transplantation results, future work could establish fully closed encapsulation-to-bioreactor culture workflows, as well as examine more tailored one-pot emulsification impeller and vessel geometries at different scales. Engineered hydrogel microenvironments could further enhance beta-cell maturation or impact performance post-transplantation. While little to no fibrosis was observed with the current 5% alginate formulation after ∼3 weeks in C57BL/6 mice in previous work, this finding should be further examined over more extended periods and in larger animal models, potentially applying chemically-modified alginates modified to reduce inflammation^52,53^. These anticipated next steps could lead to a full pipeline for upscaled cell immobilization, extended suspension culture, and transplantation of microencapsulated SC-islets.

## 4. Conclusions

SC-islets represent a promising and potentially unlimited cell source for diabetes cell replacement therapies; however, their large-scale production is hindered by agglomeration and shear-induced damage during suspension culture. In this study, we demonstrate that alginate microencapsulation provides an effective strategy to support SC-islet manufacturing and transplantation. While endocrine precursor seeding density did not significantly affect SC-islet identity, maturation, or cell recovery, differences in alginate composition were associated with shifts in endocrine cell populations, with a trend toward increased alpha-cell differentiation in stiffer environments and enhanced beta-cell maturation in softer or non-encapsulated conditions. Importantly, the emulsion-based encapsulation approach protected SC-islets during bioreactor culture, yielding high cell recovery rates during culture in a vertical wheel bioreactor, yielding high cell recovery rates over 25 days of suspension culture while preventing excessive agglomeration. Encapsulated SC-islets remained viable and functional following intraperitoneal transplantation in immunodeficient mice, secreting human C-peptide for up to 98 days and responding to physiological stimulation. Together, these findings establish alginate microencapsulation as a scalable platform for SC-islet culture and delivery and suggest that the mechanical properties of the encapsulation matrix may influence SC-islet composition and maturation.

## 5. Data availability

All data is available within the manuscript or upon request. The RNA sequencing data is available via the McGill dataverse/Borealis: https://doi.org/10.5683/SP4/WTNR06.

## 6. Acknowledgements

We thank Lisa Danielczak for supporting training, Francesco Touani Kameni and Professor. Sophie Lerouge for providing training and access to the MicroSquisher (CellScale) equipment. We thank Timothy J. Kieffer, Nicolas Proulx and Jean Ruel for insightful discussions and feedback.

This study was supported by grants from Diabetes Canada (OG-3-21-5598-CH), the Canadian Institutes of Health Research (CIHR PJT-205839 & DT1-179094), Breakthrough Type 1 Diabetes (formerly JDRF; 5-SRA-2021-1150-S-B), the Natural Sciences and Engineering Research Council of Canada (RGPIN-2020-05877), the Canadian Donation and Transplantation Research Network (CDTRP), and the Cardiometabolic Health, Diabetes and Obesity –CMDO Research Network (thematic networks supported by the FRQS, https://doi.org/10.69777/338295). A.C.R received a Recruitment Award from Biological and Biomedical Engineering department, McGill University, and McGill Engineering Doctoral Award International. J.A.B. received a Vanier scholarship from CIHR and a doctoral scholarship from the Fonds de Recherche du Québec nature et technologies (FRQNT, https://doi.org/10.69777/267872). M.B. received a doctoral scholarship from the Fonds de Recherche du Québec (FRQ, https://doi.org/10.69777/2009965).

We also acknowledge training, travel and other support provided by the following networks: the Quebec Network for Cell, Tissue and Gene Therapy –ThéCell, a thematic network supported by the Fonds de recherche du Québec–Santé (FRQS), PROTEO –The Quebec Network for Research on Protein Function (https://doi.org/10.69777/341121), CQMF/QCAM - Quebec Centre for Advanced Materials (strategic teams supported by the FRQNT, https://doi.org/10.69777/341666), the McGill Regenerative Medicine Network, the Montreal Diabetes Research Center, the Canadian Stem Cell Network, the Canadian Biomaterials Society and the Bioencapsulation Research Group.

## References

1. Cayabyab, F., Nih, L. R. & Yoshihara, E. Advances in Pancreatic Islet Transplantation Sites for the Treatment of Diabetes. Front. Endocrinol. (Lausanne). 12, (2021).

2. Kieffer, T. J., Hoesli, C. A. & Shapiro, A. M. J. Advances in Islet Transplantation and the Future of Stem Cell-Derived Islets to Treat Diabetes. Cold Spring Harb. Perspect. Med. 15, (2025).

3. Ramzy, A. et al. Implanted pluripotent stem-cell-derived pancreatic endoderm cells secrete glucose-responsive C-peptide in patients with type 1 diabetes. Cell Stem Cell 28, 2047–2061.e5 (2021).

4. Reichman, T. W. et al. Stem Cell-Derived, Fully Differentiated Islets for Type 1 Diabetes. N. Engl. J. Med. 393, 858–868 (2025).

5. Paez-Mayorga, J. et al. Emerging strategies for beta cell transplantation to treat diabetes. Trends Pharmacol. Sci. 43, 221–233 (2022).

6. Keymeulen, B. et al. Encapsulated stem cell–derived β cells exert glucose control in patients with type 1 diabetes. Nat. Biotechnol. 42, (2023).

7. Dufrane, D., Goebbels, R. M. & Gianello, P. Alginate macroencapsulation of pig islets allows correction of streptozotocin-induced diabetes in primates up to 6 months without immunosuppression. Transplantation 90, 1054–1062 (2010).

8. Vériter, S. et al. Improvement of subcutaneous bioartificial pancreas vascularization and function by coencapsulation of pig islets and mesenchymal stem cells in primates. Cell Transplant. 23, 1349–1364 (2014).

9. Thakur, G. et al. Scaffold-free 3D culturing enhance pluripotency, immunomodulatory factors, and differentiation potential of Wharton’s jelly-mesenchymal stem cells. Eur. J. Cell Biol. 101, (2022).

10. Lobel, B. T. et al. Current Challenges in Microcapsule Designs and Microencapsulation Processes: A Review. Cite This ACS Appl. Mater. Interfaces 16, 40355 (2024).

11. Lavanya, M., Namasivayam, S. K. R., Priyanka, S. & Abiraamavalli, T. Microencapsulation and nanoencapsulation of bacterial probiotics: new frontiers in Alzheimer’s disease treatment. 3 Biotech 14, (2024).

12. Lalarukh et al. Microencapsulation: An Innovative Technology in Modern Science. Polym. Adv. Technol. 36, e70066 (2025).

13. Cohen, P. J. R. et al. Engineering 3D micro-compartments for highly efficient and scale-independent expansion of human pluripotent stem cells in bioreactors. Biomaterials 295, (2023).

14. Fattahi, P. et al. Core–shell hydrogel microcapsules enable formation of human pluripotent stem cell spheroids and their cultivation in a stirred bioreactor. Sci. Rep. 11, 1–13 (2021).

15. Hoesli, C. A. et al. Pancreatic cell immobilization in alginate beads produced by emulsion and internal gelation. Biotechnol. Bioeng. 108, 424–434 (2011).

16. Hoesli, C. A. et al. Reversal of diabetes by βtC3 cells encapsulated in alginate beads generated by emulsion and internal gelation. J. Biomed. Mater. Res. - Part B Appl. Biomater. 100 B, 1017–1028 (2012).

17. Mahaddalkar, P. U. et al. Generation of pancreatic β cells from CD177+ anterior definitive endoderm. Nat. Biotechnol. 38, 1061–1072 (2020).

18. Rezania, A. et al. Reversal of diabetes with insulin-producing cells derived in vitro from human pluripotent stem cells. Nat. Biotechnol. 32, 1121–1133 (2014).

19. Balboa, D. et al. Functional, metabolic and transcriptional maturation of human pancreatic islets derived from stem cells. Nat. Biotechnol. 40, 1042–1055 (2022).

20. Iworima, D. G. et al. Metabolic switching, growth kinetics and cell yields in the scalable manufacture of stem cell-derived insulin-producing cells. Stem Cell Res. Ther. 15, 1–28 (2024).

21. Shin, D. S. et al. Mammalian cell encapsulation in monodisperse chitosan beads using microchannel emulsification. Colloids Surfaces A Physicochem. Eng. Asp. 661, 130807 (2023).

22. Brassard, J. A., Dharmaraj, S. S., Orimi, H. E. & Vdovenko, D. Iterative sacrificial 3D printing and polymer casting to create complex vascular grafts and multi-compartment bioartificial organs. (2024) doi:10.1101/2024.09.29.615298.

23. Iworima, D. G. et al. Metabolic switching, growth kinetics and cell yields in the scalable manufacture of stem cell-derived insulin-producing cells. Stem Cell Res. Ther. 15, 1 (2024).

24. Shin, D. S. et al. Mesenchymal stromal cell encapsulation in uniform chitosan beads using microchannel emulsification. 1–22 (2022).

25. Xie, Z. et al. Gene Set Knowledge Discovery with Enrichr. Curr. Protoc. 1, e90 (2021).

26. Brassard, J. A. et al. Multi-compartment bioartificial organs engineered via iterative sacrificial 3D printing and vascular integration. Cell Biomater. 0, 100455 (2026).

27. Aigha, I. I. & Abdelalim, E. M. NKX6.1 transcription factor: a crucial regulator of pancreatic β cell development, identity, and proliferation. Stem Cell Res. Ther. 11, 1–14 (2020).

28. Zhao, J. et al. PDX1+ cell budding morphogenesis in a stem cell-derived islet spheroid system. Nat. Commun. 15, 1–18 (2024).

29. Fukuda, S. et al. The intraperitoneal space is more favorable than the subcutaneous one for transplanting alginate fiber containing iPS-derived islet-like cells. Regen. Ther. 11, 65–72 (2019).

30. Prüsse, U. et al. Comparison of different technologies for alginate beads production. Chem. Pap. 62, 364–374 (2008).

31. Veiseh, O. et al. Size- and shape-dependent foreign body immune response to materials implanted in rodents and non-human primates. Nat. Mater. 14, 643–651 (2015).

32. Matlaga, B. F., Yasenchak, L. P. & Salthouse, T. N. Tissue response to implanted polymers: The significance of sample shape. J. Biomed. Mater. Res. 10, 391–397 (1976).

33. Legøy, T. A., Vet, H., Shadab, A. & Strand, B. L. Encapsulation boosts islet-cell signature in differentiating human induced pluripotent stem cells via integrin signalling. 1–16 (2020) doi:10.1038/s41598-019-57305-x.

34. Richardson, T., Kumta, P. N. & Banerjee, I. Alginate encapsulation of human embryonic stem cells to enhance directed differentiation to pancreatic islet-like cells. Tissue Eng. Part A 20, 3198–3211 (2014).

35. Ebrahim, N., Shakirova, K. & Dashinimaev, E. PDX1 is the cornerstone of pancreatic β-cell functions and identity. Front. Mol. Biosci. 9, 1–18 (2022).

36. Peterson, Q. P. et al. A method for the generation of human stem cell-derived alpha cells. Nat. Commun. 11, 1–14 (2020).

37. Iworima, D. G., Baker, R. K., Piret, J. M. & Kieffer, T. J. Analysis of the effects of bench-scale cell culture platforms and inoculum cell concentrations on PSC aggregate formation and culture. Front. Bioeng. Biotechnol. 11, 1267007 (2023).

38. Lipsitz, Y. Y., Tonge, P. D. & Zandstra, P. W. Chemically controlled aggregation of pluripotent stem cells. Biotechnol. Bioeng. 115, 2061–2066 (2018).

39. Sen, A., Kallos, M. S. & Behie, L. A. Effects of Hydrodynamics on Cultures of Mammalian Neural Stem Cell Aggregates in Suspension Bioreactors. Ind. Eng. Chem. Res. 40, 5350–5357 (2001).

40. Sart, S., Bejoy, J. & Li, Y. Characterization of 3D pluripotent stem cell aggregates and the impact of their properties on bioprocessing. Process Biochem. 59, 276–288 (2017).

41. Van Winkle, A. P., Gates, I. D. & Kallos, M. S. Mass Transfer Limitations in Embryoid Bodies during Human Embryonic Stem Cell Differentiation. Cells Tissues Organs 196, 34–47 (2012).

42. Kinney, M. A., Sargent, C. Y. & McDevitt, T. C. The multiparametric effects of hydrodynamic environments on stem cell culture. Tissue Eng. Part B. Rev. 17, 249–262 (2011).

43. Hakim, F. et al. High Oxygen Condition Facilitates the Differentiation of Mouse and Human Pluripotent Stem Cells into Pancreatic Progenitors and Insulin-producing Cells * □. 289, 9623–9638 (2014).

44. Ilc, K., Mazure, N. M., Pouysse, J., Scharfmann, R. & Duvillie, B. Oxygen Tension Regulates Pancreatic ␤ -Cell Differentiation Through Hypoxia-Inducible Factor 1 ␣. 59, (2010).

45. Nostro, M. C. et al. Stage-specific signaling through TGF b family members and WNT regulates patterning and pancreatic specification of human pluripotent stem cells. 1445, 861–871 (2011).

46. Cechin, S., et al. Influence of In Vitro and In Vivo Oxygen Modulation on b Cell Differentiation From Human Embryonic Stem Cells. 277–289 (2014).

47. Verhoeff, K. et al. Scalable Bioreactor-based Suspension Approach to Generate Stem Cell-derived Islets From Healthy Donor-derived iPSCs. Transplantation 109, e22–e35 (2025).

48. Dadheech, N. et al. Scale up manufacturing approach for production of human induced pluripotent stem cell-derived islets using Vertical Wheel® bioreactors. npj Regen. Med. 2025 101 10, 24-(2025).

49. Pollock, S. D., Galicia-Silva, I. M., Liu, M., Gruskin, Z. L. & Alvarez-Dominguez, J. R. Scalable generation of 3D pancreatic islet organoids from human pluripotent stem cells in suspension bioreactors. STAR Protoc. 4, 102580 (2023).

50. Kropp, C. et al. Impact of Feeding Strategies on the Scalable Expansion of Human Pluripotent Stem Cells in Single-Use Stirred Tank Bioreactors. Stem Cells Transl. Med. 5, 1289–1301 (2016).

51. Vegas, A. J. et al. Long-term glycemic control using polymer-encapsulated human stem cell-derived beta cells in immune-competent mice. Nat. Med. 22, 306–311 (2016).

52. Bochenek, M. A. et al. Alginate encapsulation as long-term immune protection of allogeneic pancreatic islet cells transplanted into the omental bursa of macaques. doi:10.1038/s41551-018-0275-1.

53. Vegas, A. J. et al. Corrigendum: Combinatorial hydrogel library enables identification of materials that mitigate the foreign body response in primates. Nat. Biotechnol. 34, 666 (2016).

