## Supplemental Tables and Figures for "Bridging scale-up to transplantation: pluripotent stem cell-derived pancreatic islet encapsulation in emulsion-generated high concentration alginate beads"

### Supplement

**Table 1.** Pancreatic differentiation protocol (based on previously publish protocols [17–20]

| Differentiation Stage | Lineage | Basal Media | Basal Media Supplements (Final Concentration) | Growth Factors (Final Concentration) | Small Molecules (Final Concentration) |
|---|---|---|---|---|---|
| S0D0 | Human Pluripotent Stem Cells | mTesR1 | - | - | 10 µM Y-27632 |
| S0D1 | Human Pluripotent Stem Cells | mTesR1 | - | - | - |
| S1D1-S1D3 | Definitive endoderm | MCDB 131 | 1X GlutaMAX<br>10mM D-(+)-Glucose<br>1.5 g/L Sodium Bicarbonate<br>0.5% FAF-BSA | 0.1 µg/mL Activin A Protein | 3 µM CHIR 99021 (S1D1 only) |
| S2D1-S2D3 | Primitive gut tube | MCDB 131 | 1X GlutaMAX<br>10 mM D-(+)-Glucose<br>1.5 g/L Sodium Bicarbonate<br>0.5% FAF-BSA | 50 ng/mL rh FGF-7, ACF | 1.25 µM IWP-2<br>0.25 mM L-Ascorbic acid |
| S3D1-S3D2 | Posterior Foregut | MCDB 131 | 1X GlutaMAX<br>15 mM D-(+)-Glucose<br>2.5 g/L Sodium Bicarbonate<br>2% FAF-BSA<br>0.25 mM L-Ascorbic acid<br>1X ITS-X | 50 ng/mL rh FGF-7, ACF | 0.2 µM TPB<br>0.25 µM SANT-1<br>1 µM Retinoic Acid<br>0.1 µM LDN193189 |

| Differentiation Stage | Lineage | Basal Media | Basal Media Supplements (Final Concentration) | Growth Factors (Final Concentration) | Small Molecules (Final Concentration) |
|---|---|---|---|---|---|
| S4D1-S4D4 | Pancreatic progenitor | MCDB 131 | 1X GlutaMAX<br>15 mM D-(+)-Glucose<br>2.5 g/L Sodium Bicarbonate<br>2% FAF-BSA<br>0.25 mM L-Ascorbic acid<br>1X ITS-X | 50 ng/mL rh FGF-7, ACF | 0.1 µM TPB<br>0.25 µM SANT-1<br>0.1 µM Retinoic Acid<br>0.2 µM LDN193189 |
| S4D5 (Aggrewell Plates) | Pancreatic progenitor | MCDB 131 | 1X GlutaMAX<br>15 mM D-(+)-Glucose<br>2.5 g/L Sodium Bicarbonate<br>2% FAF-BSA<br>0.25 mM L-Ascorbic acid<br>1X ITS-X | 50 ng/mL rh FGF-7, ACF | 10 µM Y-27632<br>0.1 µM TPB<br>0.25 µM SANT-1<br>0.1 µM Retinoic Acid<br>0.2 µM LDN193189 |
| S5D1-S5D5 | Endocrine progenitor | MCDB 131 | 1X GlutaMAX<br>20 mM D-(+)-Glucose<br>1.5 g/L Sodium Bicarbonate<br>2% FAF-BSA<br>1X ITS-X<br>10 µg/mL Heparin<br>10 µM Zinc Sulfate<br>100 U/mL PenStrep | - | 0.25 µM SANT-1<br>0.05 µM Retinoic Acid<br>0.1 µM LDN193189<br>0.1 µM γ-Secretase Inhibidor XX<br>10 µM Alk-5 inhibitor<br>1 µM T3 |

| Differentiation Stage | Lineage | Basal Media | Basal Media Supplements (Final Concentration) | Growth Factors (Final Concentration) | Small Molecules (Final Concentration) |
|---|---|---|---|---|---|
| S6D1-S6D7 | Immature SC-islets | MCDB 131 | 1X GlutaMAX<br>20 mM D-(+)-Glucose<br>1.5 g/L Sodium Bicarbonate<br>2% FAF-BSA<br>1X ITS-X<br>10 µg/mL Heparin<br>10 µM Zinc Sulfate<br>100 U/mL PenStrep | - | 0.1 µM LDN193189<br>0.1 µM γ-Secretase Inhibitor XX<br>10µM Alk-5 inhibidor<br>1 µM T3 |
| S7D1-D7D25 | Maturing SC-islets | MCDB 131 | 1X GlutaMAX<br>1g/L Sodium Bicarbonate<br>2% FAF-BSA<br>1X ITS-X<br>10 µg/mL Heparin<br>10µM Zinc Sulfate<br>100 U/mL PenStrep<br>1X NEAA<br>1X Trace Elements A<br>1X Trace Elements B<br>1 mM N-Acetyl-L-cysteine | - | 1 µM T3 |

**Table 2.** Catalog number and manufacturer of the basal media and supplements used in the pancreatic differentiation

| Basal Media | Basal Media Supplements | Manufacturer | Catalog # |
|---|---|---|---|
| mTesR1 | - | STEMCELL Technologies | 85850 |
| MCDB 131 | - | Gibco™ | 10372019 |
| - | GlutaMAX | Gibco™ | 35050061 |
| - | D-(+)-Glucose | Sigma Aldrich | G8769 |
| - | Sodium Bicarbonate | Thermo Fisher Scientific | S233-500 |
| - | Fatty Acid Free-Bovine Serum Albumin (FAF-BSA) | PROLIANT Health & Biologicals | 68700 |
| - | Ascorbic acid | Sigma Aldrich | A4544 |
| - | Insulin-Transferrin-Selenium-Ethanolamine (ITS-X) | Thermo Fisher Scientific | 51500056 |
| - | Heparin | Sigma Aldrich | H3149-100ku |
| - | Zinc Sulfate | Thermo Fisher Scientific | 389802500 |
| | Penicillin-Streptomycin (PenStrep) | Thermo Fisher Scientific | 15140122 |
| | Non-Essential Amino Acids (NEAA) | Gibco™ | 11140050 |
| | Trace Elements A | Corning® | 25-021-CI |
| | Trace Elements B | Corning® | 25-022-CI |
| | N-Acetyl-L-cysteine (NAAC) | Sigma Aldrich | A9165 |

**Table 3.** Catalog number and manufacturer of the growth factors used in the pancreatic differentiation

| Growth Factor or Small Molecule | Manufacturer | Catalog # |
|---|---|---|
| Y-27632 | STEMCELL Technologies | 72308 |
| Activin A Protein | Cedarlane | 338-AC-50/CF |
| CHIR 99021 | Cedarlane | 4423/10 |
| rh FGF-7, ACF | Thermo Fisher Scientific | 78186.2 |
| IWP-2 | Cedarlane | 13951 |
| Ascorbic acid | Sigma Aldrich | A4544 |
| TPB | Sigma Aldrich | 565740 |
| SANT-1 | Cedarlane | 14933 |
| Retinoic Acid | Sigma Aldrich | R2625 |
| LDN193189 | Sigma Aldrich | SML0559 |
| Secretase Inhibitor XX | Sigma Aldrich | 565789 |
| Alk-5 inhibitor | Cedarlane | 14794 |
| 3,3′,5-Triiodo-L-thyronine sodium salt (T3) | Sigma Aldrich | T6397 |

**Table 4.** Conjugated antibodies used in the Flow Cytometry analysis across various pancreatic differentiation stages

| Pancreatic Differentiation Stage | Lineage | Antibody | Vendor & Catalog # |
|---|---|---|---|
| Stage 1 Day 3 | Definitive endoderm | SOX17-PE | BD Biosciences/561591 |
| | | PE mouse IgG1 | BD Biosciences/554680 |
| Stage 4 Day 3 | Pancreatic progenitor | PDX1-PE | BD Biosciences/562161 |
| | | NKX6.1-AF647 | BD Biosciences/563338 |
| | | NEUROD1-PE | BD Biosciences/563001 |
| | | PE mouse IgG1 | BD Biosciences/554680 |
| | | AlexaFluor® 647 mouse IgG1 | BD Biosciences/557732 |
| Stage 7 Day 10 & 25 | Maturing SC-islets | PDX1-PE | BD Biosciences/562161 |
| | | NKX6.1-AF647 | BD Biosciences/563338 |
| | | NEUROD1-PE | BD Biosciences/563001 |
| | | Glucagon-PE | BD Biosciences/565860 |
| | | C-peptide-AF647 | BD Biosciences/565831 |
| | | PE mouse IgG1 | BD Biosciences/554680 |
| | | AlexaFluor® 647 mouse IgG1 | BD Biosciences/557732 |

**Table 5.** Gene Symbols and Full Gene Names for Differentially Expressed Genes Identified by RNA Sequencing

| Gene Symbol | Full Gene Name |
|---|---|
| CPA2 | Carboxypeptidase A2 |
| CTRB2 | Chymotrypsinogen B2 |
| FXYD2 | Carboxypeptidase A2 |
| AJAP1 | Chymotrypsinogen B2 |
| FXYD2 | FXYD Domain Containing Ion Transport Regulator 2 |
| AJAP1 | Adherens Junctions Associated Protein 1 |
| CAMK2A | Calcium/Calmodulin Dependent Protein Kinase II Alpha |
| LAMA3 | Laminin Subunit Alpha 3 |
| COL2A1 | Collagen Type II Alpha 1 Chain |
| SLC2A2 | Solute Carrier Family 2 Member 2 (GLUT2) |
| SIK1 | Salt Inducible Kinase 1 |
| MMP9 | Matrix Metallopeptidase 9 |
| MMP13 | Matrix Metallopeptidase 13 |
| EZR | Ezrin |
| PDLIM3 | PDZ And LIM Domain 3 |
| MYADM | Myeloid Associated Differentiation Marker |
| ANGPT2 | Angiopoietin 2 |
| AHNAK | AHNAK Nucleoprotein |
| F3 | Coagulation Factor III (Tissue Factor) |
| ADCYAP1 | Adenylate Cyclase Activating Polypeptide 1 (PACAP) |

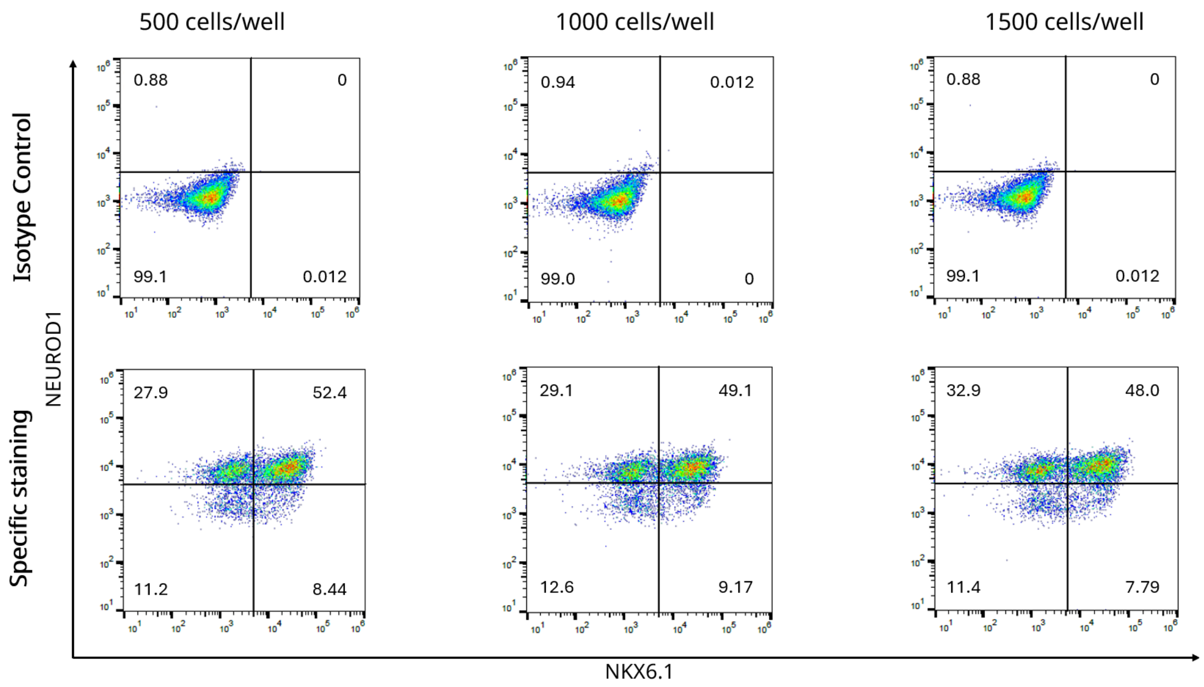


**Figure S1** – Representative flow cytometry graphs showing S7D10 population expression of NEUROD1 vs NKX6.1.

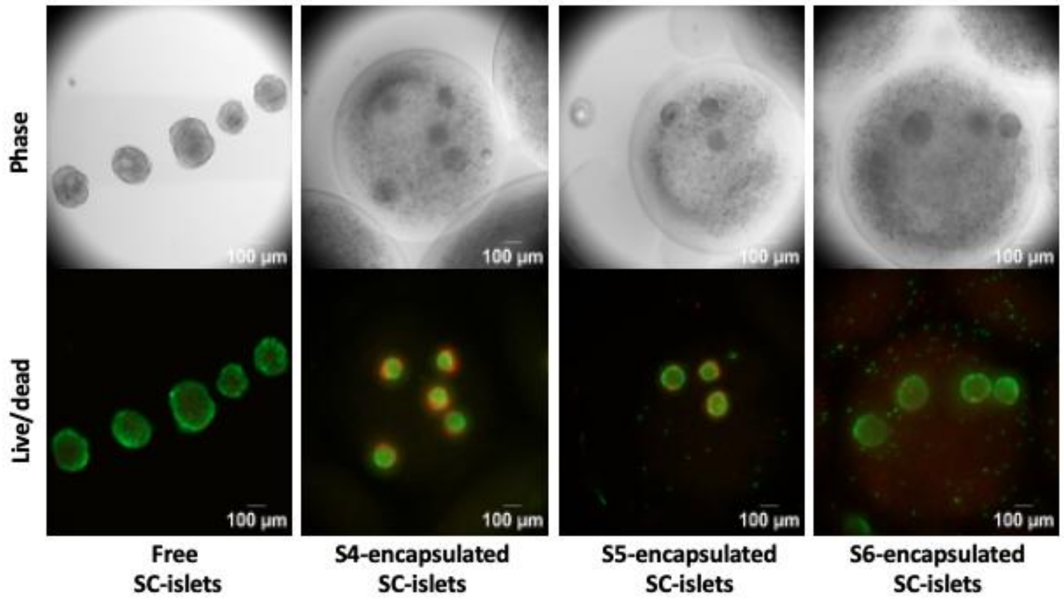


**Figure S2** – Impact of differentiation stage on viability of stem cell-derived islets after 24h. Representative phase-contrast images (top row) and Live/Dead staining (bottom row) of free and encapsulated SC-islets at different stages (S4, S5, S6).

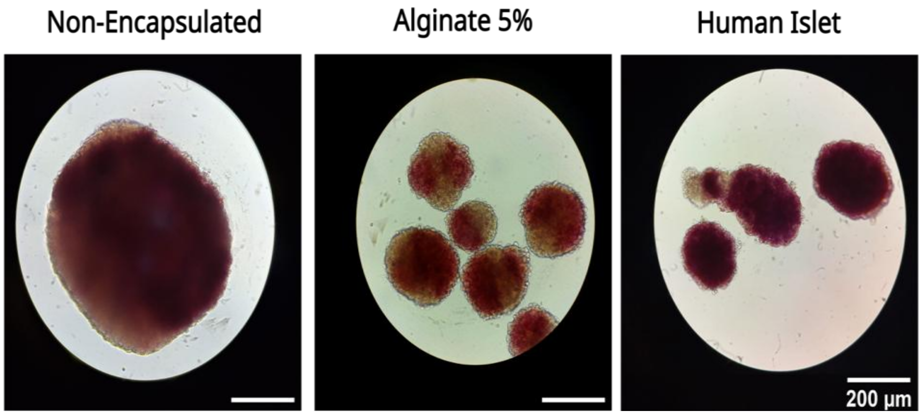


**Figure S3** - Dithizone staining of maturing SC-islets, non encapsulated, and encapsulated after 3 weeks of culture in the PBS mini bioreactor (aggregates recovered from degelled beads) (N=2). Human islets were used as a control

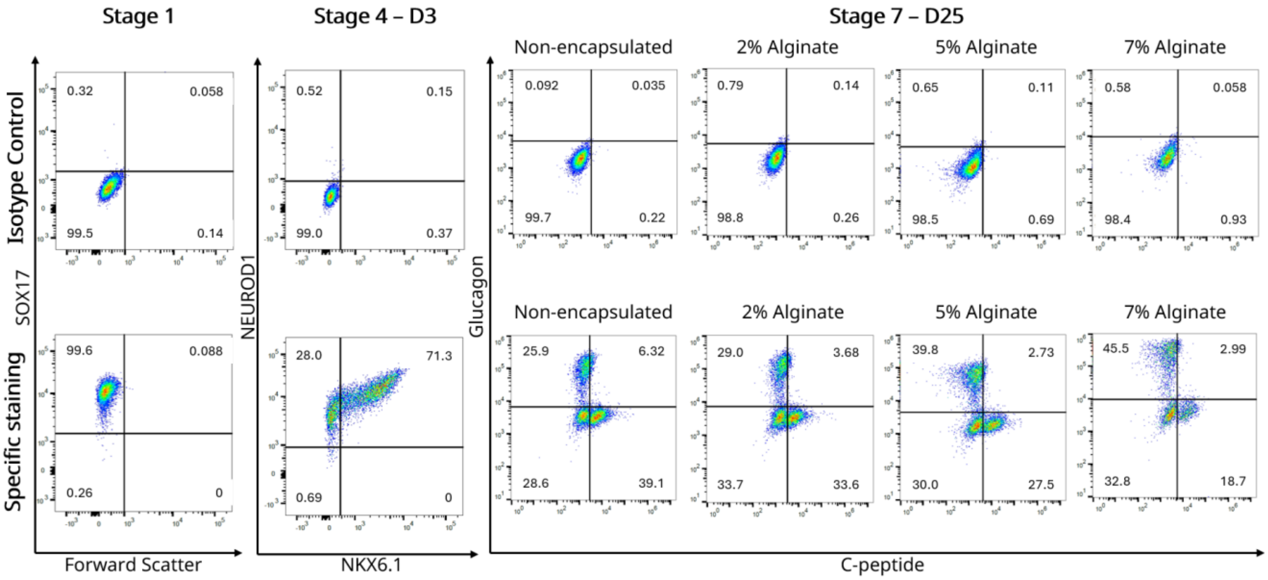


**Figure S4 –** Representative flow cytometry graphs showing stage 7-day 25 population expression of genes of interest during the stem cell-derived pancreatic islet differentiation protocol.
